# A millisecond coarse-grained simulation approach to decipher allosteric cannabinoid binding at the glycine receptor *α*1

**DOI:** 10.1101/2023.04.19.537578

**Authors:** Alessio Bartocci, Andrea Grazzi, Nour Awad, Pierre-Jean Corringer, Paulo C.T. Souza, Marco Cecchini

## Abstract

Glycine receptors (GlyR) are regulated by small-molecule binding at several allosteric sites. Cannabinoids like tetrahydrocannabinol (THC) and N-arachidonyl-ethanol-amide (AEA) potentiate GlyR but their mechanism of action is not fully established. By combining millisecond coarse-grained MD simulations powered by Martini 3 with backmapping to all-atom representations, we characterize the cannabinoid-binding sites at zebrafish GlyR-*α*1 with atomic resolution. Based on hundreds of thousand ligand-binding events, we find that cannabinoids bind to the transmembrane domain of the receptor at both intrasubunit and intersubunit sites. For THC, the intrasubunit binding mode predicted in simulation is in excellent agreement with recent cryo-EM structures, while intersubunit binding recapitulates in full previous mutagenesis experiments. Intriguingly, AEA is predicted to bind at the same intersubunit site despite the strikingly different chemistry. Statistical analyses of the receptor-ligand interactions highlight potentially relevant residues for GlyR potentiation, offering experimentally testable predictions. The predictions for AEA are validated by electrophysiology recordings of rationally designed mutants. The results highlight the existence of multiple cannabinoid-binding sites for the allosteric regulation of GlyR and put forward an effective strategy for the identification and structural characterization of allosteric sites in transmembrane proteins.

## Introduction

Glycine receptors (GlyR) are pentameric ligand-gated ion channels that play a critical role in motor coordination and essential sensory functions such as vision and audition. ^1^ They are integral transmembrane protein assemblies featuring a large extracellular domain (ECD) that hosts two or more glycine-binding sites and a compact transmembrane domain (TMD) that forms an axial chloride channel through the postsynaptic membrane. These receptors mediate synaptic inhibition in the spinal cord and the brain stem ^2^ and have since long been recognized as pharmacological targets for hyperekplexia, temporal lobe epilepsy, and more recently chronic pain.^3^ At the structural level, GlyR is by far the best characterized pentameric ligand-gated ion channel with more than 40 high-resolution structures solved in different conformations (i.e., resting, pre-active, active, and desensitized) and in complex with modulatory ligands such as agonists, partial agonists, antagonists and allosteric modulators.^4–8^ A wide panel of small-molecule compounds including psychoactive drugs, general anesthetics, and neurotoxins are known to regulate the GlyR function.^3,9^ Recently, a library of 218 unique chemical entities with documented modulatory activity at homomeric GlyR-*α*1 and GlyR-*α*3 along with a structural annotation of their binding site on the receptor has been collected (GRALL).^10^ Strikingly, about one third of it appears to target the TMD, thus highlighting the relevance of this region for the allosteric modulation of synaptic receptors. Cannabis is the most commonly used psychoactive drug worldwide.^11^ Phytocannabinoids like Δ^9^-tetrahydrocannabinol (THC) and cannabidiol (CBD), i.e., the primary psychoactive compounds in cannabis, are lipid-like signaling molecules that were shown to modulate the glycinergic response in addition to targeting the cannabinoid receptors CB1 and CB2.^12^ The endocannabinoid N-arachidonyl-ethanol-amide (AEA), a.k.a. anandamide, was also found to potentiate the glycinergic response in oocytes expressing recombinant GlyR-*α*1.^13^ And THC, CBD and other exogenous cannabinoids, i.e., ajulemic acid, HU-210, and WIN 55212-2, were found to potentiate GlyR non-competitively.^14^ Despite its pharmacological relevance, the molecular mechanism of GlyR modulation by cannabinoids has remained elusive, mostly due to lack of structural information at high resolution.

Functional studies by patch-clamp electrophysiology in combination with site-directed mutagenesis have shown that the non-conservative mutation of serine 267 to iso-leucine on the transmembrane helix M2 at human GlyR-*α*1 abolishes co-activation by CBD, ajulemic acid and HU-210.^15^ And serine substitution at position 296 on the transmembrane helix M3 of purified human GlyR-*α*3 was shown to abolish potentiation by CBD.^16^ Most recently, Kumar et al. reported cryo-EM structures of zebrafish GlyR-*α*1 (Zf-GlyR-*α*1) reconstituted in lipid nanodiscs in complex with THC.^17^ These structures suggest that THC binds to a lipid-exposed intrasubunit pocket at the interface of the transmembrane helices M3 and M4. However, the intrasubunit nature of this site, which is inconsistent with the topographical location of known allosteric sites at GlyR and pentameric homologues, ^18^ as well as the existence of unassigned lipid-like densities compatible with additional THC-binding sites ^17^ indicate that further studies are needed to establish the mechanism of the THC-induced potentiation of GlyR. Moreover, the allosteric modulation by endocannabinoids like AEA remains to be explored.

Most drugs currently on the market have been designed to target the primary active site of proteins, a.k.a. the orthosteric site. Allosteric ligands that bind to topographically distinct sites offer a competitive advantage over orthosteric compounds as they are more selective, they limit the risk of off-target effects, and can be used synergistically with known drugs to potentiate or attenuate the pharmacological response.^19^ Moreover, allosteric sites have been recently exploited to develop therapeutics for proteins that were considered undruggable. ^20^ Nonetheless, the identification of allosteric sites in proteins remains challenging and their discovery has mostly been serendipitous.^21^ Systematic approaches for the identification and structural characterization of allosteric modulatory sites need to be developed, particularly for transmembrane proteins.

The use of coarse-grained (CG) models preserving chemical specificity such as the Martini force field^22^ has emerged as a powerful and cost-effective strategy to probe the spatial and temporal evolution of biomolecules,^23–25^ particularly for pharmacological applications.^26,27^ This modeling approach provides a smoother potential energy surface that allows for integration time steps up to 30 fs in molecular dynamics (MD) simulations.^28^ The latter yields a speed-up of 2-3 orders of magnitude relative to all-atom MD, which opens to the exploration of time scales and sizes that were previously out of reach. Recently, the newly parameterized version of Martini, a.k.a. Martini 3, ^29,30^ in combination with unbiased MD was shown to reproduce the experimental binding modes and binding affinity of several small molecules to a protein with high accuracy (*<* 0.5 kcal/mol) and no a priori knowledge of the ligand-binding site.^31^ In addition, the expansion of the bead chemical types and sizes in Martini 3 allows for a better coverage of the chemical space, which facilitates CG modeling of small-molecule drugs.^32,33^ Last, refinement of the ligand binding poses by back-mapping to all-atom representations^34,35^ opens to multi-scale simulation approaches to explore protein-ligand binding at equilibrium.

A fast-growing area for Martini simulations is the analysis of protein–ligand interactions in the transmembrane region of pharmacologically relevant targets. Several attempts have been reported that aim at the identification and structural characterisation of ligand-binding sites,^36–38^ ranking of binding modes,^31,39,40^ and structural refinement of the protein/ligand complex via all-atom backmapping.^34^ Since the parameterisation of new drug-like compounds in Martini remains challenging, these studies are often limited to the exploration of protein-lipids interactions,^41^ while CG explorations of modulatory ligands (e.g. agonists, antagonists and allosteric modulators) has remained rare and of qualitative nature.^27^

Here, we combine efficient CG simulations powered by Martini 3 with backmapping to all-atom resolution to decipher the allosteric binding site(s) of two cannabinoids, i.e., THC-Δ^9^ and AEA, at the active state of Zf-GlyR-*α*1. Our strategy allows for mapping of the cannabinoid-receptor interaction in the membrane environment at equilibrium, it provides direct probing of the cannabinoid binding/unbinding kinetics, and opens to the structural characterization of statistically relevant binding modes with atomic resolution. Based on hundred of thousands ligand-binding events, we find that THC and AEA bind at both intrasubunit and intersubunit sites in the transmembrane domain of the receptor. The simulations reveal the existence of a previously uncharacterized intersubunit cannabinoid-binding site, which complements recent structural biology data and recapitulates in full mutagenesis studies on the potentiation by THC. Predictions on AEA are validated by electrophysiological recordings of rationally designed mutants, providing the first characterization of endocannabinoid binding to GlyR-*α*1. Taken together, the results highlight the existence of multiple, topographically distinct cannabinoid-binding sites for the allosteric modulation of synaptic receptors and put forward an effective simulation approach for their identification and structural characterization.

## Results

### GlyR-THC recognition

The interaction between THC and the active state of Zf-GlyR-*α*1 was explored by 0.5 millisecond, unbiased, coarse-grained molecular dynamics (CG/MD) simulations with 5% THC in the lipid membrane; see Methods for details. These simulations were used to sample the receptor-cannabinoid interaction in the membrane at equilibrium; see Supplementary Movie 1. The spatial distribution of THC around the transmembrane domain of the GlyR active state is shown in Figure 1A. The density maps in the upper and lower leaflets show a highly symmetric distribution with little or no THC density in the interior of the protein, including the ion pore (white region). The highly homogeneous color at a distance from the protein-lipid interface (light-blue region) and the symmetric distribution of the binding hot-spots (dark blue) indicate converged sampling of the protein-ligand interaction. In addition, the bright color of the hot-spots indicates that THC binds at these sites specifically, which suggests the existence of cannabinoid-recognition sites. When the THC density is analysed in 3D, the data reveal a density peak in the mid of the membrane bilayer in proximity of the transmembrane helices M3 and M4 from the same subunit (i.e. intrasubunit); see Figure 1B. Moreover, additional density is found between subunits at the interface of the M3 (−) and M1 (+) helices. Altogether, the results indicate that the CG/MD simulations provide converged sampling of the GlyR-THC interaction in the lipid membrane and highlight the presence of one or more recognition sites for THC in the active state of the receptor.

**Figure 1:**
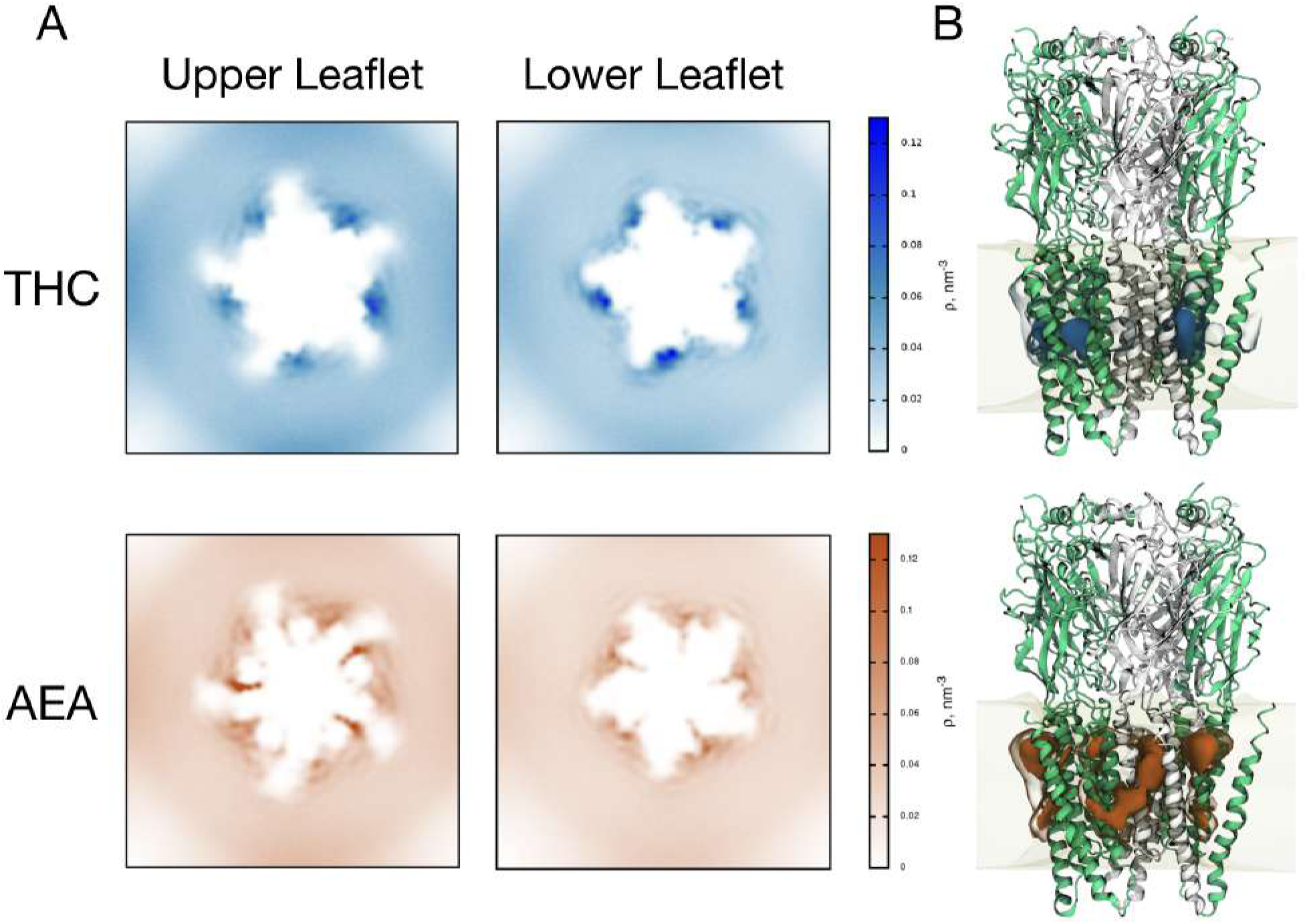
Equilibrium distribution of cannabinoid ligands around GlyR. (A) 2D-density maps (in nm^−3^) for THC (blue) and AEA (orange) in the lipid membrane split in upper and lower leaflets as shown in Supplementary Fig. 1. Note that the lower-density zones at the corners of the simulation box result from the re-centering the MD trajectory on the protein rather than a sampling issue. (B) 3D-density maps. The protein subunits are color-coded in green and white alternately to highlight the interfaces and distinguish between intrasubunit and intersubunit binding. The membrane environment is shown as a semi-transparent region. In all cases, darker colors correspond to binding hot-spots.

In order to characterize the GlyR-THC interaction and isolate specific recognition events, the simulation trajectories were analyzed by monitoring the number of protein-ligand contacts over time (see Methods). Using a dual cut-off scheme to reduce the statistical noise, 91266 THC-binding events were collected along with the corresponding residence times; see Supplementary Method 4. With the help of Eq. 1, the statistics collected over 0.5 ms simulation at 300 K and 5% THC for the bound and the unbound states of the ligand reveal a standard free energy of binding of −3.613 ± 0.003 kcal/mol; see Table 1. In the limit of the accuracy of the CG parameterisation, this result indicates that THC binds rather weakly to the GlyR active state with a *K_d_* of 2.3 mM. Nonetheless, the probability for the protein to be bound is remarkably high (98.1%), which is due to both the high initial concentration of THC in the simulations (i.e., 0.011 M) and the 6-fold increase in its effective concentration due to complete partitioning in the membrane; see Table 1. Remarkably, the use of Eq. 1 in combination with ligand-binding probabilities extracted from hundreds of thousands of binding events yields an estimate of the standard free energy of binding with a statistical uncertainty *<* 0.01 kcal/mol. These results demonstrate that the CG/MD approach provides straightforward access to protein-ligand binding affinities (*K_d_*) in the membrane with impressive statistical precision; see Supplementary Method 3.

**Table 1:** Binding affinity predictions from CG/MD simulations.

| System | N | V<br>(nm<sup>3</sup>) | C<br>([M]) | V<sub>memb</sub><br>(nm<sup>3</sup>) | C<sub>memb</sub><br>([M]) | $p_b$ | $p_u$ | $\Delta G_{\text{bind}}^\circ$<br>(kcal/mol) | $K_d$<br>([mM]) |
| --- | --- | --- | --- | --- | --- | --- | --- | --- | --- |
| THC | 16 | 2328 | 0.011 | 397 | 0.067 | 0.234 | 0.766 | $-3.613 \pm 0.003$ | 2.3 |
| AEA | 16 | 2326 | 0.011 | 400 | 0.066 | 0.299 | 0.701 | $-3.811 \pm 0.008$ | 1.7 |

In addition to the thermodynamics, CG/MD simulations open to direct estimates of the binding kinetics (*K_off_* ). The distribution of the residence time of THC on GlyR (Supplementary Fig. 15) indicates that 98.6% of the binding events sampled by CG/MD are short lived and correspond to non-specific interactions at the protein-lipid interface. However, a small but significant fraction of them (i.e. 1290 events) were long-lived (*>* 100ns) and represent receptor-ligand recognition events; see Supplementary Table 6. Upon clustering binding events by RMSD and correcting for the pentameric symmetry of the receptor (see Methods), a collection of 255 unique THC-binding modes were extracted. Their probability distribution in Figure 2A highlights the existence of three dominant binding modes that account altogether for ∼40% of GlyR-THC recognition sampled in simulation. Analysis of the residence times per binding mode reveals monoexponential distributions with a characteristic time of 160 ns for the most probable one (orange), and 78 ns and 66 ns for the second (red) and third (cyan); (Figure 2C). Interestingly, the longest-lived binding mode is also the most populated one.

**Figure 2:**
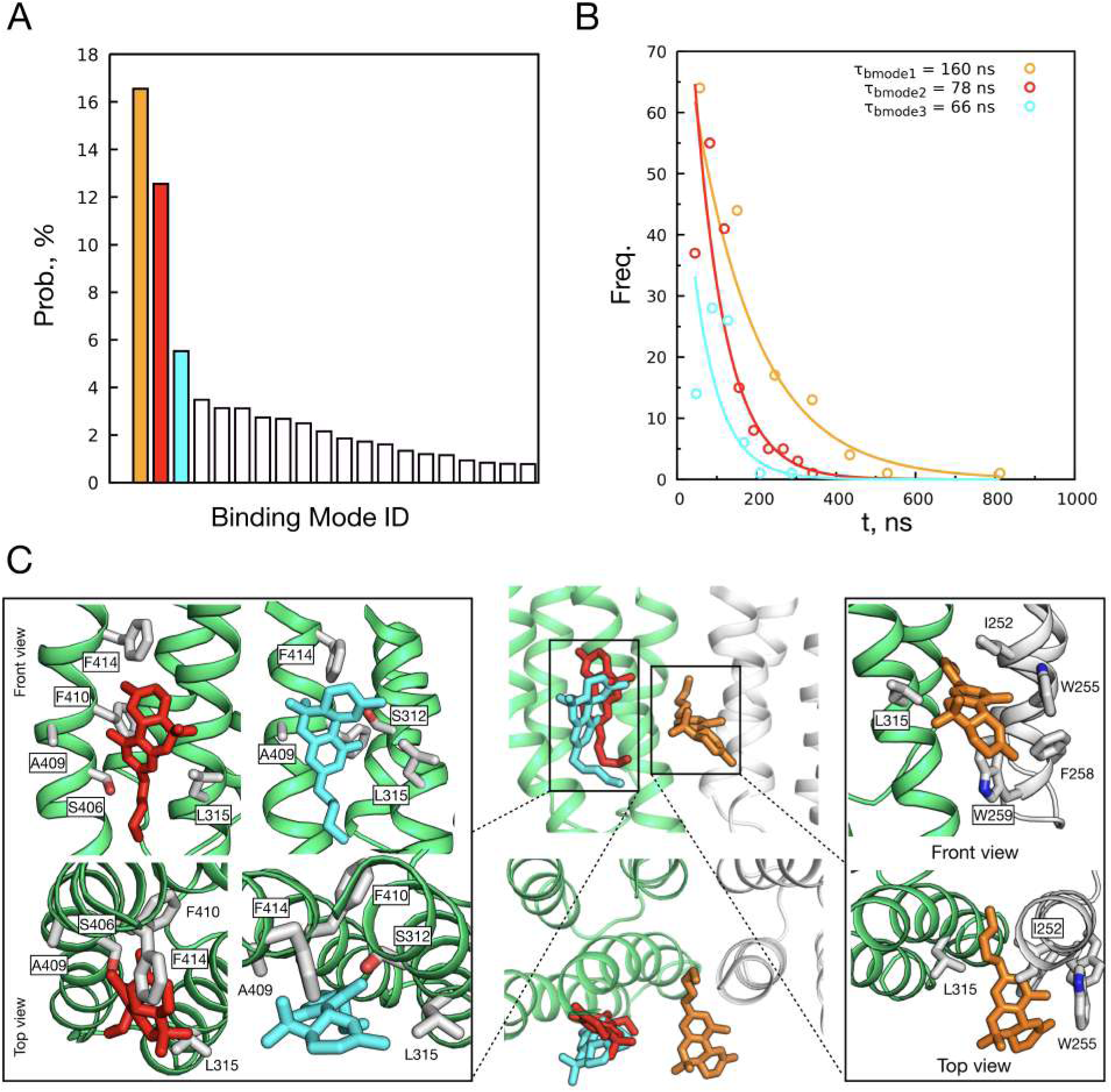
THC allosteric binding at GlyR. (A) Specific binding modes are represented with associated probabilities. The dominant binding modes are color-coded. (B) Residence time (*τ_r_*) distributions of the three most important binding events. The characteristic residence time per binding mode is given. The fit was done as described in Methods. (C) Atomistic representation of the dominant binding modes. Protein chains are color-coded in green and white to highlight adjacent subunits. The THC molecule is shown in sticks and colored following the scheme in panel A. For each binding mode, the protein residues involved in the receptor-ligand recognition are highlighted in the insets.

Upon backmapping to an all-atom representation, the structural models of the three dominant binding modes reveal that THC may bind specifically both intersubunit and intrasubunit. When THC binds intersubunit, its terpenic core is sandwiched between two tryptophans from helix M1 (W255 and W259), and its short alkyl tail inserts in a hydrophobic pocket at the interface of the M3 (−) and M1 (+) helices (orange, Figure 2C). When THC binds intrasubunit (red or cyan), the ligand positions at the interface between M3 and M4 and forms a H-bond with a nearby serine. Notably, the two intrasubunit binding modes target the same pocket but involve flipping of THC by 180°, which re-positions its H-bonding moiety relative to residues that line the pocket. Consistent with the density plots in Figure 1, the all-atom representations of the three dominant binding modes by THC correspond to binding hot-spots. Despite being less populated, the intrasubunit binding modes correspond to the darkest spots in Figure 1. Our analysis reveals that these hot spots actually account for two binding modes (red and cyan in Figure 2C), which artificially increases the statistical weight of intrasubunit binding.

In order to pinpoint residues that could be relevant for GlyR potentiation by THC, a receptor-ligand contact analysis was carried out on the simulation frames corresponding to the three dominant binding modes; see Supplementary Fig. 19 and Methods for details. The results predict that five residues in the TMD are critical for binding THC intersubunit, with W255, F258 and W259 from M1 stabilizing the terpenic core of the ligand via stacking interactions, and L315 from M3(−) and I252 from helix M1(+) forming a hydrophobic pocket where the short alkyl chain of THC slips in. In addition to those, six residues are relevant for binding THC intrasubunit, with S312 or S406 forming a H-bond with the phenol group of the ligand, and A409, F410, and F414 from M4 and L315 from M3 forming a hydrophobic pocket that accommodates its terpenic core. A recent mutagenesis analysis by Kumar et al^17^ revealed that residue substitutions at W255, F258, W259, S312 and F410 significantly decrease or even abolish GlyR potentiation by THC; for residue numbering consider a +8 shift in the sequence of Kumar (7M6O) relative to the sequence used here (6PM6). Strikingly, all residues by Kumar were identified as critical for binding THC by our CG/MD simulation analysis (Supplementary Fig. 19). Moreover, the literature observation that serine substitution at position 296 (i.e. S312 in 6PM6) abolishes GlyR potentiation by CBD^16^ is also consistent with predictions from our simulations. These comparisons show that the statistical analysis of the receptor-THC interactions sampled by CG/MD simulations at equilibrium is remarkably consistent with available mutagenesis data.

### GlyR-AEA recognition

A similar investigation based on millisecond CG/MD simulations was carried out to characterize the interaction between GlyR and anandamide (AEA). The results show a different interaction pattern relative to THC, with AEA penetrating much deeper in the TMD and being mostly localized intrasubunit in the upper leaflet and intersubunit in the lower leaflet (Figure 1). Similar to THC, the probability of finding AEA at the receptor-lipid interface is higher than the bulk at more than one interaction site.

A detailed analysis of the receptor-ligand contacts reveals that 129346 binding events were sampled by 0.5 ms CG/MD at 300 K and 5% of AEA in the membrane, 1.5 % of which had a residence time longer than 100 ns and represents specific recognition events; see Supplementary Table 6. The distribution of the residence time reveals that AEA binding to the GlyR active state is longer-lived than THC, with binding events lasting up to 6 *µs*, which is six times longer than the longest lived THC-binding event (Figure 3). Similar to THC, the simulations indicate that AEA binding to the GlyR active state is weak, albeit slightly more favorable than THC, with a 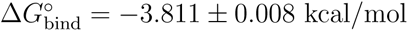, which corresponds to a *K_d_* of 1.7 mM and an increase in affinity of 0.2 kcal/mol relative to THC; see Table 1. Statistics over the long-lived binding events highlight the existence of four dominant binding modes among 192 sampled in total, which account for more than 60 % of the specific GlyR-AEA recognition (Figure 3A). Two of them (green and cyan) have a residence time *τ_r_ >* 800 ns, whereas the other two (red and yellow) are shorter-lived with residence times *τ_r_* ≃ 200-300 ns (Figure 3B). Upon backmapping to all-atom representations, the dominant binding modes reveal that the two longest-lived modes correspond to intrasubunit binding at an upper leaflet site (green and cyan, Figure 3C), whereas the two shorter-lived binding modes correspond to binding to upper leaflet (orange) or lower leaflet (red) intersubunit sites.

**Figure 3:**
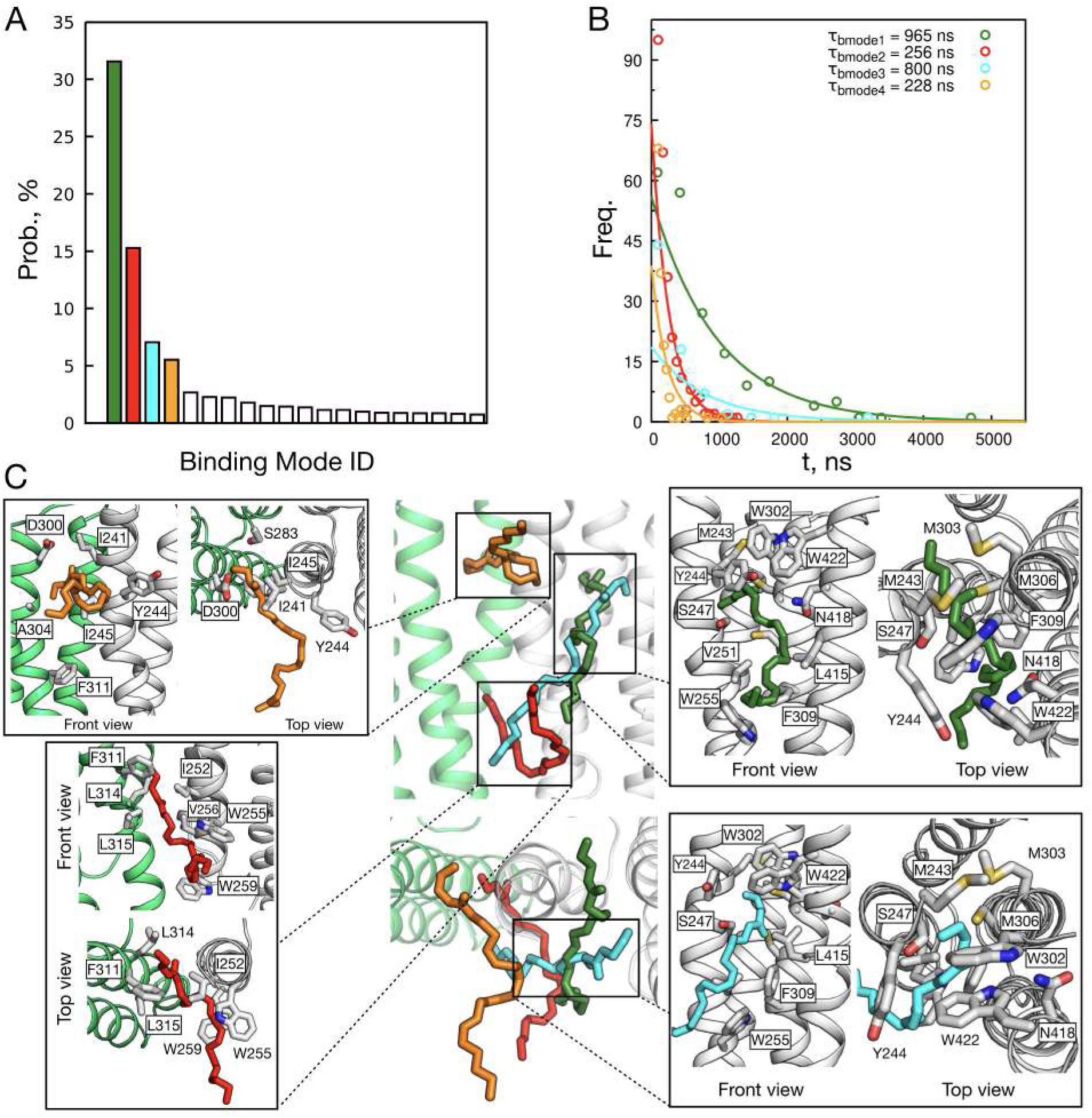
AEA allosteric binding at GlyR. (A) Specific binding modes are represented with associated probabilities. The dominant binding modes are color-coded. (B) Residence time (*τ_r_*) distributions of the four most important binding events. The characteristic residence time per binding mode is given. The fit was done as described in Methods. (C) Atomistic representation of GlyR-AEA interaction in the dominant binding modes. Protein chains are color-coded in green and white to highlighting adjacent subunits, while AEA molecule is shown in sticks and color-coded following the scheme in panel A. For each binding mode, the protein residues involved in the receptor-ligand recognition are highlighted in the insets.

To identify residues that could be potentially relevant for GlyR potentiation by AEA, a statistical analysis of the per-residue receptor-ligand contacts was carried out over 0.5 ms of CG/MD simulation (Figure 4). The results reveal that: i) binding modes 1 and 3 are essentially equivalent (Supplementary Fig. 17); ii) when AEA binds intrasubunit, the ligand inserts in a hydrophobic cavity at the interface of the transmembrane helices M1, M3, and M4 and anchors its head via polar contacts with the side chains of S247 from helix M1 and C306 from helix M3, while the rest of the molecule makes non-polar contacts with the side chains of M243, Y244, Y251 and W255 from M1, W302, M303 and F309 from M3, and L415, N418 and W422 from helix M4; iii) when AEA binds intersubunit at the upper leaflet site (orange), the ligand inserts the polar head in the cleft formed by the M3(+) and M1(−) helices and establishes a H-bond with the side chain of S283 from helix M2 similar to ivermectin binding,^4,5^ while forming non-polar contacts with two isoleucines (I241 and I245) and one tyrosine (Y244) from helix M1 and one alanine (A304) from helix M3; iv) when AEA binds intersubunit to the lower leaflet site, molecular recognition involves the surprising insertion of the AEA polar head in the same hydrophobic pocket occupied by the alkylic tail of THC in its most populated binding mode along with extensive interactions formed with W255 and W259 from M1 that sandwich part of the alkylic tail of the ligand. We note that in the lower intersubunit binding mode (red) the polar head of AEA inserts deeper than THC and was found to establish polar interactions with water molecules percolating from the ion pore via several water channels; see Supplementary Fig. 18. Remarkably, the CG/MD simulations predict that the same residues implicated in THC binding (i.e. I252, W255, W259, and L315) are critical for AEA binding, despite the strikingly different chemistry of these two cannabinoid ligands. In addition, they highlight the implication of S283 (S267 in human) in AEA recognition, whose role in GlyR’s potentiation by cannabinoids or ivermectin has been previously recognized in functional assays^15^ and cryo-EM studies, ^4,5^ respectively.

**Figure 4:**
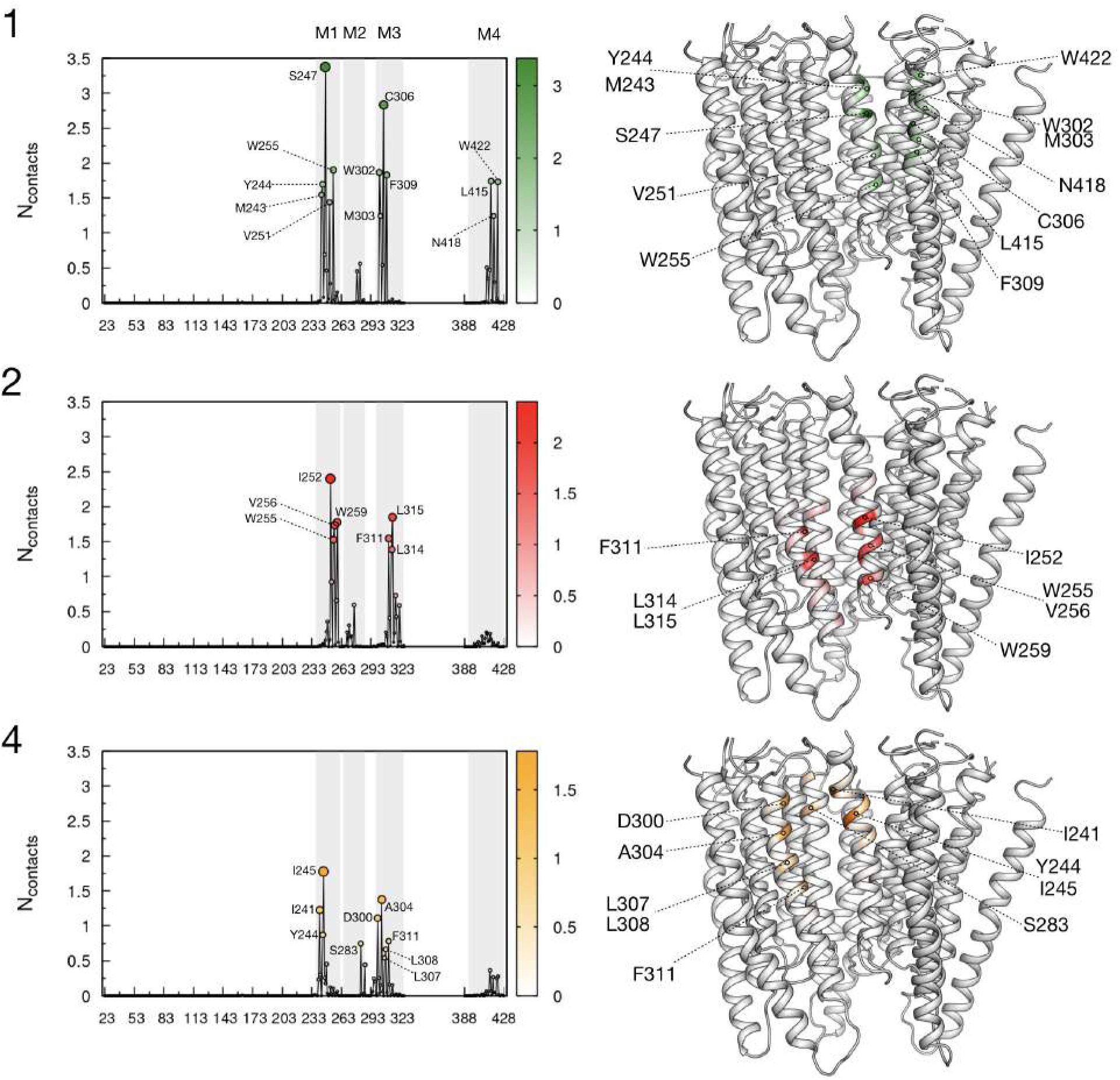
GlyR-AEA contact analysis. The analysis was carried out for the three dominant GlyR-AEA binding modes predicted by CG/MD (Figure 3, green, red and orange) as described in Methods. On the left-hand side, the average number of contacts per residue is plotted along the sequence of the protein. Gray boxes indicate protein stretches corresponding to the transmembrane helices M1-M4. On the right-hand side, the residues identified by the contact analysis are indicated on the protein structure. The color code is the same as that in Figure 3.

### Mutagenesis studies on GlyR potentiation by AEA

The contact analysis based on converged CG/MD simulations highlight residues that are mostly involved in AEA recognition at the TMD of Zf-GlyR-*α*1 (Figure 3). The modeling results indicate that: i. S247 (M1), C306 (M3), and W422 (M4) are involved in binding AEA intrasubunit; ii. I252, W255, V256, and W259 on M1 of the (−)-subunit and F311, L314, and L315 on M3 of the (+)-subunit are involved in binding AEA at the lower inter-subunit site; and iii. I241 and I245 on M1 of the (−)-subunit, A304 on M3 of the (+)-subunit and S283 on M2 of the (+)-subunit are involved in binding AEA at the upper inter-subunit site; here residue numbering corresponds to zebrafish GlyR-*α*1 in 6PM6. By focusing on inter-subunit binding and prioritizing residues previously tested for THC modulation,^17^ a full electrophysiological characterization of seven mutants at the lower intersubunit site (hW239, hS241, hF242, hW243, hF295, hL298, and hL299 corresponding to W255, S257, F258, W259, F311, L314, and L315 in 6PM6) and three mutants at the upper intersubunit site (hI225, hI229, and hS267 corresponding to I241, I245, and S283 in 6PM6) of human GlyR-*α*1 was carried out to explore their effect on the allosteric modulation by AEA; see Supplementary Fig. 21 and Methods for details. In addition, two mutations that were previously reported to affect GlyR potentiation by AEA, i.e., hS296A was shown to abolish potentiation^16^ and hK385A to attenuate it to 14%,^14^ were also explored. Following the protocol of Kumar et al, ^17^ phenylalanine, leucine, isoleucine and serine residues were mutated into alanine, whereas the bulkier tryptophans were replaced by phenylalanines. Dose-response curves of the twelve mutants showed significant deviations from the wild type (WT), particularly for five loss-of-function mutations, i.e., hW239F, hS241A, hF242A, hW243F, and hL298A, see Supplementary Fig. 22. Therefore, the effect of residue substitution on AEA potentiation was analysed by comparing gating currents elicited by an EC20 concentration of glycine in presence of 10 *µ*M AEA. The results show that co-application of glycine and AEA produces a 1.5-fold potentiation at human GlyR-*α*1 and that eight out of twelve mutants tested display strong deviations from WT. Strikingly, alanine substitution at hS241, hF295, hL298, and hL299 completely abolishes AEA potentiation, whereas alanine substitution at hI225, hS296, and hF242 decreases it. Lastly, the replacement of hW243 by phenylalanine increases AEA potentiation by 50%. Surprisingly and unlike previously reported,^14^ no effect of alanine substitution at K385 on the outer pre-M4 helix was detected. When located on the cryo-EM structure of Zf-GlyR-*α*1 solved in complex with THC,^17^ the eight residues having a significant effect on AEA potentiation highlight three topographically distinct regions in the TMD of the receptor (Figure 5B). The first group gathers five residues at both M1(−) and M3(+) helices, whose mutations have highly significant effects on the GlyR potentiation (hS241, hW243, hF295, hL298, hL299) and clearly highlight the lower intersubunit binding site identified by CG/MD (red). The second group involves hI225 and suggests the implication of residues at the upper intersubunit site (orange), particularly at the water/membrane interface. The third group includes hS296 (white) and provides evidence that the intrasubunit site targeted by THC^17^ is also involved in the potentiation by AEA. Altogether, the mutagenesis experiments demonstrate a primary role of the lower intersubunit site on the potentiation of GlyR by AEA. In addition, they highlight the existence of multiple and topographically distinct ligand-binding sites for the allosteric modulation of GlyR-*α*1 by cannabinoids.

**Figure 5:**
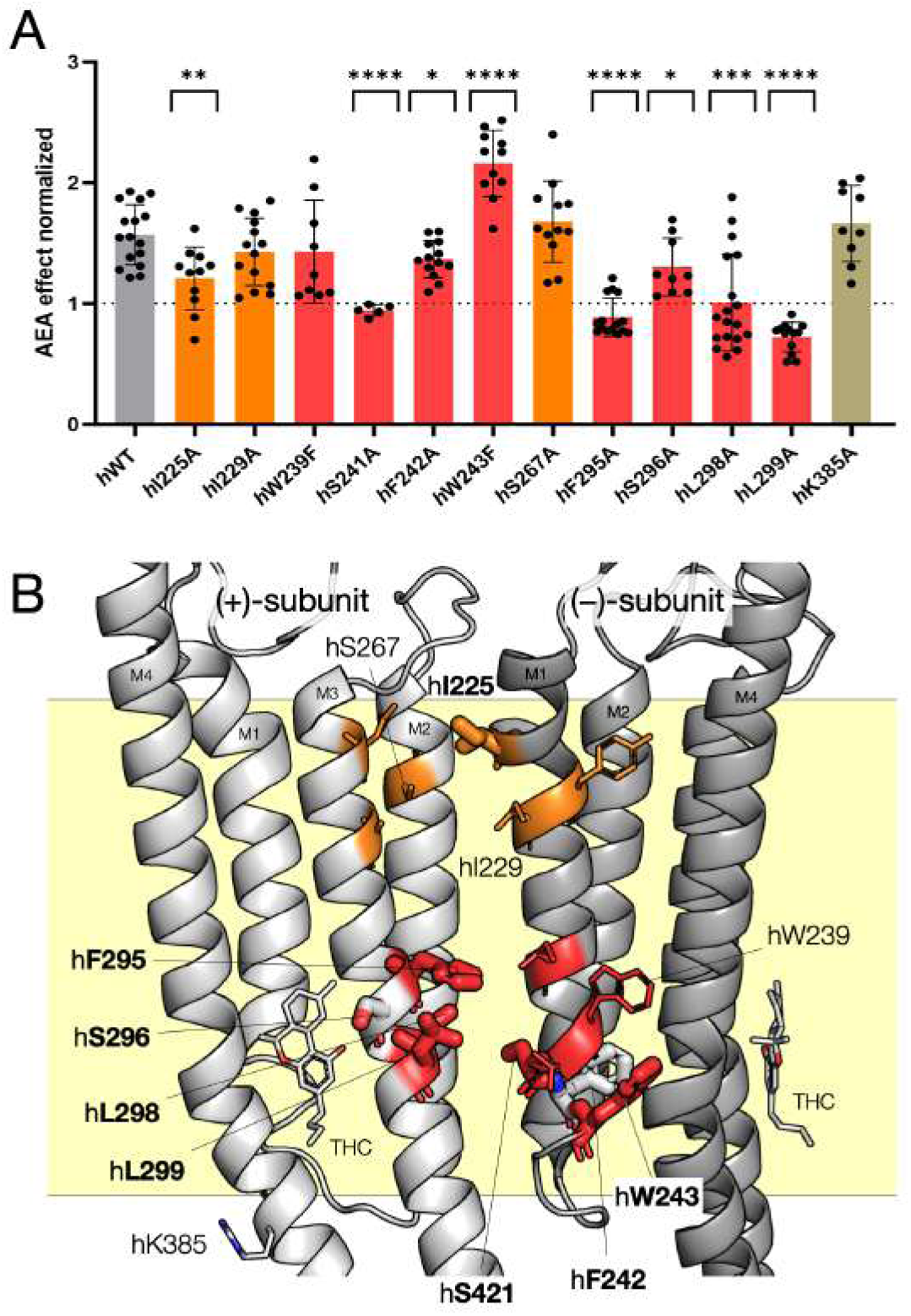
Mutational analysis to characterize the AEA-binding site(s) at human GlyR-*α*1. (A) AEA potentiation (as peak of AEA-glycine current/peak glycine current) is shown for WT (n=16), hS267A (n=12), hI225A (n=11), hS296A (n=9), hW239F (n=9), hS241A (n=5), hI229A (n=14), hK385A (n=9), hF242A (n=13), hW243F (n=11), hF295A (n=15), hL298A (n=18), hL299A (n=13). Data are shown as mean ± s.d. for (n) independent experiments. Electrophysiology experiments were performed on independent oocytes, from multiple different surgeries. Two-sided Mann-Whitney test ^**^*P* = 0.0039 (hI225A),^*^*P* = 0.0196 (hS296A), ^****^*P* ≤ 0.0001 (hS241A), ^*^*P* = 0.0401(hF242A), ^****^*P* ≤ 0.0001 (hW243F), ^****^*P* ≤ 0.0001 (hF295A), ^***^*P* = 0.0003 (hL298A), ^****^*P* ≤ 0.0001 (hL299A). (B) The structural location of residues explored by mutational studies is highlighted on the cryo-EM structure of zebrafish GlyR-*α*1 solved in complex with THC.^17^ Amino acids represented as thick sticks correspond to residues whose substitution yields a significant effect on the potentiation by AEA in electrophysiology. The red and orange colors correspond, respectively, to amino acids at the lower and upper intersubunit binding sites as identified by the CG/MD simulations. THC molecules and hS296 lining the intrasubunit allosteric site implicated in THC potentiation are shown in white.

## Discussion

Endocannabinoids like anandamide (AEA) and phytocannabinoids like tetrahydrocannabinol (THC) regulate the function of synaptic receptors including the ionotropic glycine receptor (GlyR).^1^ Due to the lack of structural information at high resolution, the molecular mechanism of GlyR potentiation by endo- and phytocannabinoids is not fully established. Recently, cryoEM structures of GlyR-*α*1 from zebrafish reconstituted in lipid nanodiscs illuminated the receptor in complex with THC.^17^ These structures highlighted the existence of an intrasubunit cannabinoid-binding site located in the transmembrane domain of the protein at the interface of the M3 and M4 helices. The intrasubunit nature of this ligand-binding site is surprising and inconsistent with structural biology of GlyR and pentameric homologues, which shows that allosteric modulatory sites are typically located at the subunit-subunit interface.^10,42^ Moreover, the different chemistry of THC and AEA prevents from a straight-forward extrapolation of the structural results for THC, so that the nature of the allosteric modulatory site targeted by endocannabinoids remains to be elucidated.

We set out to provide an atomistic representation of the GlyR-cannabinoid interaction using a multi-scale simulation approach based on the synergistic combination of efficient coarse-grained MD simulations powered by Martini 3, which open to the millisecond timescale on commodity computer resources, with backmapping to all-atom representations to explore the details of molecular recognition with atomic resolution. Based on hundreds of thousand ligand-binding events, the simulations reveal the existence of multiple cannabinoid-binding sites in the transmembrane domain of GlyR and provide atomistic representations of the dominant binding modes. As a bonus, the simulations provide converged results for the ligand-binding affinity (*K_d_*) and residence time (*K_off_*), which are useful to explore the thermodynamics and kinetics of the cannabinoid-receptor interaction in the limit of the approximations of the energy model in use. The results for THC reveal the existence of two allosteric sites for phytocannabinoids that account for ∼40% of the specific receptor-ligand recognition sampled in simulation. One of them is located intrasubunit and lies at the interface between the transmembrane helices M3 and M4. The other one is intersubunit and is located between helix M3 from the principal subunit and helix M1 from the complementary subunit. Strikingly, the intrasubunit allosteric site predicted in simulation perfectly matches the THC-binding mode illuminated by the cryoEM structures of Kumar et al^17^ (Figure 6A), which validates the modeling approach. In addition, the intersubunit binding mode overlaps with the neurosteroid-binding mode illuminated by the X-ray structure of GLIC-GABA*_A_*R chimera in complex with tetrahydrodeoxycorticosterone (PDB ID: 5OSB)^43^ (Figure 6B) and is consistent with unassigned electron density present in the cryo-EM maps of Kumar et al;^17^ see Supplementary Fig. 20. Intriguingly, the discovery of an intersubunit binding site for THC offers a plausible re-interpretation of recent site-directed mutagenesis experiments showing that residue substitution (to alanine or phenyl-alanine) at W255, P266 and F410 (6PM6 numbering) abolishes GlyR potentiation by THC without changing the sensitivity to glycine;^17^ note that these residues correspond to W263, P274, and F418 in Kumar’s numbering (7M6O) due to a +8 residue shift relative to 6PM6. To explain the data, the existence of an allosteric coupling between the intrasubunit THC-binding site and the desensitization gate of GlyR (P266) via a network of aromatic residues involving the M1-M2 linker was postulated.^17^ The existence of an intersubunit THC-binding site at the M1-M3 interface as predicted by our CG/MD simulations offers a more natural explanation of the mutagenesis data and provides further support to our investigation.

**Figure 6:**
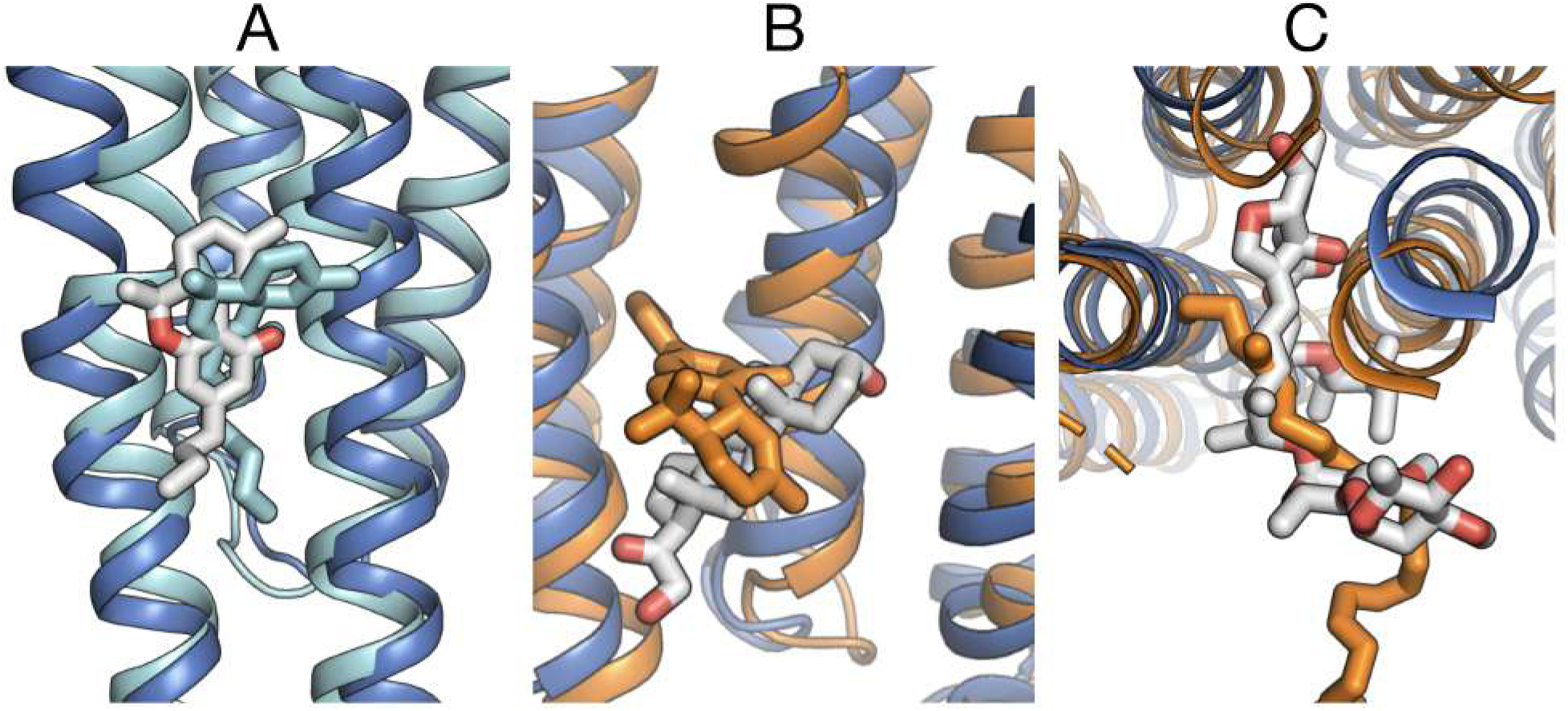
Comparison of cannabinoid binding modes predicted by CG/MD with experiments. (A) Superimposition of the cryo-EM structure of zebrafish GlyR-*α*1 solved in complex with THC (PDB ID: 7M6O) with the intrasubunit binding mode for THC predicted in simulation (Figure 2, cyan). The heavy-atom RMSD between experimental and predicted ligand-binding modes is 2.7 Å. (B) Superimposition of X-ray structure of the GLIC-GABA*_A_*R chimera solved in complex with tetrahydrodeoxycorticosterone (PDB ID: 5OSB) with the THC intersubunit binding mode predicted in simulation (Figure 2, orange). (C) Superimposition of the X-ray structure of human GlyR-*α*_3_ solved in complex with ivermectin (PDB ID: 5VDH) with the AEA upper intersubunit binding mode predicted in simulation (Figure 3, orange).

The simulation results on AEA predict that fatty-acid endocannabinoids and di-terpene phytocannabinoids bind to the same lower intersubunit allosteric site. Also, they show that, unlike THC, AEA may bind to an upper intersubunit pocket that partly overlaps with the binding site of the positive allosteric modulator ivermectin (PDB ID: 5VDH)^5^ (Figure 6C). Since THC and AEA display similar modes of action on the glycinergic response^14^ despite the strikingly different chemistry, the existence of a common lower intersubunit binding site as predicted by CG/MD (i.e. same binding pocket and comparable binding kinetics) offers a plausible mechanism for the allosteric modulation of GlyR by endocannabinoids. Moreover, the simulation analysis provides precise experimentally testable predictions. In fact, if AEA-binding to the lower intersubunit site is involved in GlyR’s potentiation, the mutagenesis analysis of Kumar et al^17^ with AEA should show a significant reduction (if not abolition) of the potentiation in some of those mutants. Vice versa, if AEA-binding to the upper intersubunit site were most involved in the potentiation mechanism, the serine to iso-leucine non-conservative mutation of S283 (S267 in human), which was shown to abolish potentiation by cannabidiol, ajulemic acid and HU-210, ^15^ would have a detectable effect in presence of AEA. By capitalizing on the simulation results, ten mutants (i.e., seven at the lower intersubunit site and three at the upper intersubunit site) along with two other extracted from the literature (h296A^16^ and hK385A^14^) were expressed and their potentiation by AEA characterized by two-electrode voltage-clamp (TEVC) electrophysiology. Strikingly, seven out of ten mutations predicted in silico displayed highly significant effects on the potentiation by AEA ranging from considerable attenuation to full abolition (Figure 5). These mutagenesis results thus demonstrate a primary implication of the lower intersubunit site on the potentiation mechanism by AEA and highlight the existence of other (at least two) topographically distinct binding sites that are relevant for the allosteric modulation of GlyR by cannabinoids. These results provide a structural characterization of the allosteric modulatory site(s) for endocannabinoids at GlyR-*α*1.

More generally, the combination of high-resolution structures with CG/MD simulations and functional studies with site-directed mutagenesis put forward an effective strategy for the identification and structural characterization of allosteric sites in transmembrane proteins. Starting with high-resolution structures of the protein, e.g. from cryo-EM, CG/MD simulations powered by Martini 3 are broadly accessible and straightforward to run.^44,45^ The striking efficiency of these calculations yields converged sampling of the configurational space, which provides atomistic representations of the statistically relevant binding modes along with a list of residues involved in protein-ligand recognition. A characterization of the CG/MD predictions by electrophysiology in combination with site-directed mutagenesis then follows to provide an experimental validation and consolidate the conclusions. Using two cannabinoids known to potentiate GlyR-*α*1 (i.e., THC and AEA), we have shown that this strategy is remarkably efficient. Considering that the TMD of the receptor without the intracellular domain contains 134 residues, the synergistic combination of CG/MD simulations with electrophysiology and mutagenesis allowed for the characterization of one new allosteric site in the transmembrane domain at 7% investment relative to an exhaustive search (i.e., 10 mutations over 134 residues), which would be even lower in case of heteropentameric receptors. In addition, our results shown that the combination of efficient CG/MD simulations with robust backmapping harnesses the complexity beyond protein-ligand recognition in lipid membranes, which results from the competition of multiple ligands including lipids at topographically distinct sites in a highly crowded environment, remarkably well. The same is currently out of reach by standard modeling approaches and possibly structural biology too. In fact, while all-atom MD and docking are deemed to fail due to incomplete sampling and/or inappropriate ranking of ligand-binding modes, the structural studies are likely to fail because of their tendency to provide coordinates for a single binding mode, which might be only partly relevant for the allosteric modulation particularly in the transmembrane region. On the other hand, ligand parameterisation in Martini 3 is challenging and remains the bottleneck of the simulation protocol presented here. The latter would require the establishment of robust and efficient strategies for an automatic parameterisation of ligands in Martini 3, which have recently started to emerge. ^46^

In conclusion, the results presented in this paper provide fundamental insights on the cannabinoid-GlyR interaction. This work highlights an unexpected complexity underlying ligand binding and allosteric regulation at synaptic receptors and presents an effective simulation approach to harness it. The striking reproduction of the cryo-EM binding mode of THC to zebrafish GlyR-*α*1 as well as the rational design of mutants strongly affecting GlyR potentiation by AEA demonstrate that coarse-grained MD simulations have come to age and will be a useful tool for the identification and structural characterization of allosteric transmembrane sites with little or no a priori knowledge.

## Methods

### Setup for the CG/MD simulations

Classical coarse-grained, molecular dynamics (CG/MD) simulations were performed using GROMACS version 2021^47^ with the Martini 3 force field. ^29,31^ Initial coordinates of zebrafish GlyR-*α*1 in its open, ion-conducting state were obtained from the PDB (6PM6).^8^ The protein was mapped to a CG representation using martinize 2.0.^48–50^ An elastic network with a distance cutoff of 0.8 nm was applied on the backbone to preserve the pentameric symmetry of the protein. The force constant of the elastic network was set to 1500 *kJ mol*^−1^ *nm*^−2^, which yields an RMSD of the backbone beads of 2.4 Å from the initial coordinates after 10 *µ*s MD without freezing the transmembrane helices; see Supplementary Method 1.

The protein model was embedded in a 1-palmitoyl-2-oleoyl-*sn*-glycero-3-phosphocholine (POPC) bilayer (164/173 in the upper/lower leaflets) with 5% (16 copies) of THC or AEA using insane.py^51^ (Figure 7A). Using the same tool, the system was solvated with water, Cl^−^ counterions, and 0.15 M concentration of NaCl. CG parameters for tetrahydrocannabinol (THC) and N-arachidonyl-ethanol-amide (AEA) were derived as described below.

**Figure 7:**
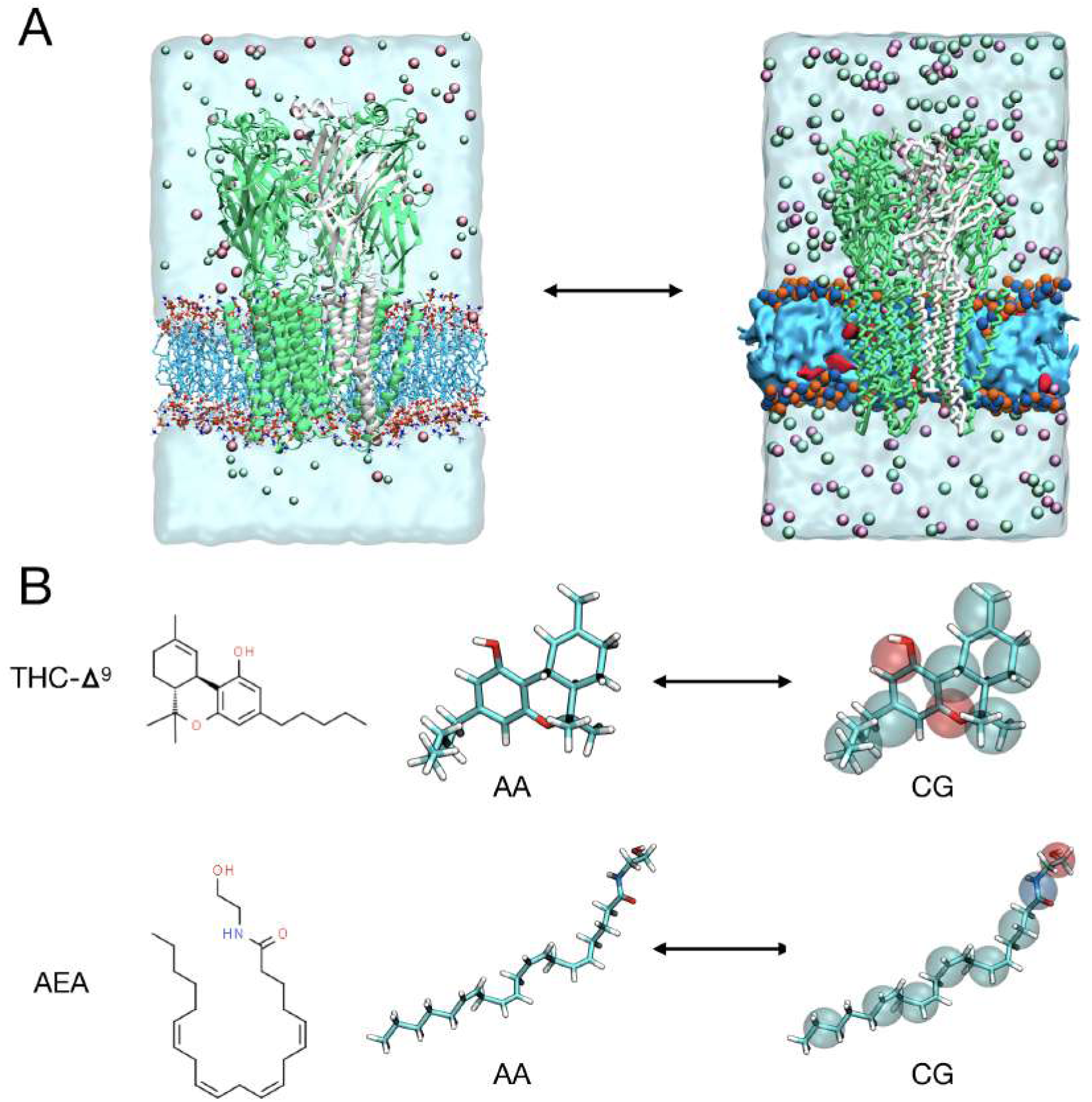
CG/MD simulation setup. (A) The GlyR active state (PDB ID: 6PM6) was embedded in a POPC lipid membrane with 5% of cannabinoids. The protein is shown as white and green cartoons. The POPC polar heads are shown as blue and orange spheres with the alkylic tails represented as a cyan surface. The cannabinoid ligands are shown in red. Ions are shown as green (Na^+^) and violet (Cl^−^) spheres, while water molecules are represented by a light-blue continuum. (B) On the left, the chemical structure of tetrahydrocannabinol (THC) and N-arachidonyl-ethanol-amide (AEA) are shown. All-atom (AA) representations of the two cannabinoids are given in the middle. Corresponding coarse-grained (CG) representations are given on the right with beads represented as semi-transparent colored spheres.

### Ligand parameterization

Initial coordinates for (−)-Δ^9^-tetrahydrocannabinol (THC) and N-arachidonyl-ethanol-amide (AEA) were obtained from the GRALL database^18^ and converted into CHARMM36-FF^52,53^ topologies using the CGENFF python-tool.^54^ CG parameters for the ligands were obtained following standard procedures in Martini 3^30^ as described in Supplementary Method 2. The CG mappings are presented in Supplementary Fig. 2. For the bonded interactions, CG parameters were validated by comparing bond distances, angles and dihedrals as well as molecular volume and SASA with all-atom MD results for the same molecules in solution. Parameters for the non-bonded interactions were validated by comparing water/octanol partition free energies of the CG models with log *P_oct/wat_* values from the literature; see Supplementary Table 2. For the reference simulations in all-atom MD, THC and AEA were simulated with 4000 TIP3P^55^ water molecules in the NPT ensemble at 298K and 1 bar.

### MD simulations

For the CG/MD simulations, the molecular systems were energy minimized for 50000 steps using the steepest descent algorithm and then equilibrated at room temperature for 100 ns with position restraints on the protein backbone (1000 *kJ mol*^−1^ *nm*^−2^ force constant) and 1 *µs* without restraints. Production runs of 10 *µs* were carried out in 50 replicates with different initial velocities for a total sampling of 0.5 ms per molecular system. A 20 fs integration time step *t_step_*was used and periodic boundary conditions applied. During the equilibration and production stage, the temperature was maintained constant at 300 K using a modified Berendsen thermostat^56^ with a *τ_T_* coupling constant of 1 ps. The pressure was maintained at 1 bar (semiisotropic coupling) using a Berendsen barostat^57^ during the initial equilibration (*τ_P_* = 4 *ps*, compressibility of 3 · 10^−4^ *bar*^−1^) and a Parrinello-Rahman barostat^58^ (*τ_P_* = 12 *ps*, compressibility of 3 · 10^−4^ *bar*^−1^) for the second equilibration and all production runs. Electrostatic interactions were cutoff at 1.1 nm using the reaction-field method,^59^ while Lennard-Jones interactions were cutoff at 1.1 nm using the potential-shift Verlet method. Bonds between beads were constrained to equilibrium values using the LINCS algorithm.^60^ During the 1 *µs* MD equilibration and all production runs, an elastic network acting on the protein backbone was used to preserve the quaternary structure of the protein.

### MD analysis tools

#### Binding affinity calculations

Protein-ligand binding affinities or *K_d_* were accessed in simulation using the following expression of the standard free energy of binding

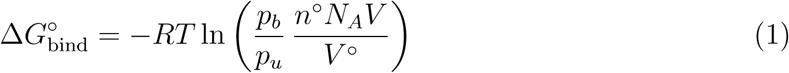

with *p_b_* and *p_u_* being, respectively, the probability for the ligand to be bound or unbound, *V*the solution volume, *N_A_* the Avogadro constant, and *n*^◦^ and *V* ^◦^ the number of moles and the solution volume at standard conditions, i.e., typically 1 *mol/L* concentration; see Supplementary Method 3 for the derivation of Eq. 1 and Ref.^61^ for an alternative derivation. This expression provides straightforward access to *K_d_* values from converged CG/MD simulations and was shown to be independent of the initial concentration of the ligand as well as the number of ligand copies in the simulation box; see Supplementary Method 3. In this work, *p_b_* and *p_u_* were estimated via a protein-ligand contacts analysis of the simulated trajectories with a distance cutoff of 0.6 nm. Specifically, all simulation frames with a number of contacts larger than one were considered as representatives of the bound state, whereas those with zero contacts as representatives of the unbound state. The solution volume, *V* , in Eq. 1 was determined as the volume of the simulation box minus the excluded volume of the protein quantified by the Voss Volume Voxelator software^62^ via the 3V webserver; see Supplementary Method 3 for details.

The statistical uncertainty on the standard free energy of binding was estimated as

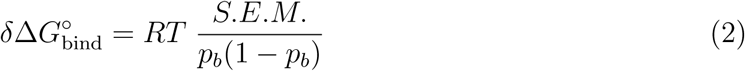

with *S.E.M.* being the standard error of the mean for *p_b_*that was estimated from multiple simulation runs of 10 *µ*s; see Supplementary Method 3 for the derivation of Eq. 2.

### Density maps

2D density maps were generated using the gmx densmap tool. The upper and lower membrane leaflets were defined as the volume slices (along the axis perpendicular to the membrane) between the center of mass of the membrane and the POPC heads (NC3 and PO4 beads); see Supplementary Fig. 1. The 3D densities maps were obtained by computing the occupancy of the ligands in the three-dimensional space using the volmap plugin of vmd,^63^ with the protein aligned in the center of the box. The grid points had a distance of 2 Å. All ligand beads were taken into account in their actual size; i.e. the radius of the regular and small beads were 2.64 Å and 2.30 Å, respectively, while that of the tiny beads was 1.91 Å.

### Binding modes and residence time

Ligand-binding events were identified by monitoring the number of protein-ligand contacts using a distance cutoff of 0.6 nm. To reduce the statistical noise on the determination of their time duration, a dual-cutoff scheme on the number of contacts was used. For this purpose, 10 contacts (cutoff *_bound_*) were used to set the beginning of a binding event and 5 contacts (cutoff *_unbound_*) for the unbinding (Supplementary Fig. 14). Binding events lasting longer than 100 ns were annotated as specific, extracted from the MD trajectories and analyzed; they represent approximately 1% of the total for both THC and AEA; see Supplementary Fig. 15 and Supplementary Table 6. Representative structures per binding event were obtained by RMSD clustering of ligand coordinates using the gmx cluster tool and using the center of the most populated cluster. To correct for the pentameric symmetry of the receptor, representative structures were extracted after alignment of the protein to a common subunit-subunit interface. Upon sorting binding events by time duration, representative structures were grouped into binding modes using a leader clustering algorithm with an RMSD cutoff of 0.34 nm starting from the longest-lived binding event as reference; the cutoff value of 0.34 nm was fine-tuned by trials and errors. The probability per binding mode was then determined by counting the total number of frames associated to it. For THC, the entire molecule was considered for the RMSD calculation, for AEA only the head beads were considered. The residence time per binding mode was obtained by fitting the distribution of the time duration per mode using a single exponential law as 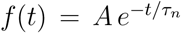 with *τ_n_* being the residence time of the n-*th* binding mode. Last, atomistic representations of the most populated binding modes were obtained using backward.^34^ For this purpose, GlyR, POPC, THC/AEA and ions were described with the CHARMM36 force field,^52,53^ while water with the TIP3P^55^ model. The backmap files to produce all-atom representations of the ligands were generated from scratch.

### Critical residues for binding

To identify residues primarily involved in cannabinoid binding to GlyR, a protein-ligand contact analysis was carried out on the ensemble of snapshots corresponding to the dominant binding modes. For this purpose, the gmx mindist tool was used to compute the average number of protein-ligand contacts per residue (N *_contacts_*) using a bead distance cutoff of 0.5 nm. By plotting *N*_contacts_ along the sequence of the protein per binding mode, the most relevant residues naturally emerge; see Figure 4.

### Experimental section

#### DNA cloning and mutagenesis

cDNA constructs for human GlyR-*α*1 and all associated point mutations in this study were cloned into the pMT3 vector. The mutations were introduced using site-directed mutagenesis protocol.

#### Electrophysiological recordings by two-electrode voltage-clamp (TEVC) in oocytes

Xenopus laevis oocytes were obtained from EcoCyte Bioscience, Germany. Oocytes were injected with cDNA (at 10-30 ng/*µ*l) encoding for human GlyR-*α*1 and mutants and incubated at 19^◦^C in Barth’s solution containing: 88 mM NaCl, 1 mM KCl, 0.66 mM Na(NO_3_)_2_, 0.75 mM CaCl_2_, 0.82 mM MgSO_4_, 2.4 mM NaHCO_3_, 10 mM HEPES, pH adjusted to 7.6 with NaOH. Oocytes expressing receptors were recorded 2-4 days after injection. Currents were recorded using a Warner OC-725C amplifier, and a Digidata 1550 A interface. Currents were analyzed by Clampfit 10.7.0 (Molecular Devices). Oocytes were clamped at a holding potential of −60 mV and solutions were exchanged using a syringe pump perfusion system flowing at a rate of 12 ml/min. The electrophysiological solutions consisted of (in mM) 96 NaCl, 2 KCl, 1.8 CaCl_2_, 1 MgCl_2_, and 5 HEPES, pH adjusted to 7.6 with NaOH. Glycine was purchased from Sigma-Aldrich and anandamide (AEA) was purchased from Enzo Life Sciences. AEA was co-perfused with glycine at the concentrations described in the text, following dilutions of at least 1000-fold from stock concentrations in either distilled water or DMSO. Data and statistical analysis were performed using Graphpad Prism 10.1 (Graphpad Software). All data are reported as mean + s.d. for (n) individual oocytes. The dose-response parameters (*EC*_50_ and Hill coefficients, *nH*) were obtained from curve fitting of normalized concentration-response data-points to the equation *I*_Gly_ = *I*_min_ + (*I*_max_ − *I*_min_)[Gly]*^nH^/*([Gly]*^nH^* + *EC*_50_*^nH^*). The maximal current (*I*_max_) corresponded to the average maximal current elicited by a concentration of 5 mM glycine. All statistical tests were unpaired and two-sided.

## Data Availability

Initial coordinates for the protein were obtained from the RCSB Protein Data Bank under the accession code 6PM6. Initial coordinates for the two cannabinoid ligands were obtained from GRALL. For structural comparisons the following PDB structures were used: 5OSB, 5VDH, 7M6O, 7M6M. Coarse-grained models, Martini 3 parameters, and GROMACS input files (itp and gro files) for THC and AEA are freely available from the MAD webserver. ^45^ Molecular snapshots representative of the most populated ligand-binding modes generated in this study along with their backmapped atomistic representations have been deposited in Zenodo [https://doi.org/10.5281/zenodo.12206040]. Source data are provided with this paper. The raw data underlying the binding affinity calculations for THC and AEA and all model calculations reported in the Supplementary Information are provided in the Source Data file. A summary table to explicitly direct readers to information regarding reproducibility of the MD simulations is provided as Supplementary Fig. 23.

## Acknowledgments

This work received funding from grants of the Agence nationale de la Recherche (ANR–18– CE11–0015) to M.C. and P.J.C. and the European Union’s Horizon 2020 Research and Innovation Programme under Marie Skolodowska-Curie Grant Agreement 956314 [ALLODD] to M.C. and ERC (Grant no. 788974, Dynacotine) to P.J.C. Support from the French National Center for Scientific Research (CNRS) and funding from a research collaboration agreement with PharmCADD to P.C.T.S. are gratefully acknowledged. Computational resources and support at the HPC centers of the University of Strasbourg (Mesocentre) and the University of Reims (ROMEO) are gratefully acknowledged. P.C.T.S. acknowledges the support of the Centre Blaise Pascal’s IT test platform at ENS de Lyon (Lyon, France) for the computer facilities. The platform operates the SIDUS solution developed by Emmanuel Quemener. ^64^

## Author Contributions

MC conceived the study and obtained funding. AB and MC designed the simulation protocol with contributions from PCTS. AB and PCTS parameterized the ligands and protein coarsegrained models. AB performed the MD simulations. AG and MC developed the theory to extract ligand-binding affinities from MD simulations with contributions from PCTS. AB wrote the ligand occupancy and residence time analysis tools, while AG the binding free energy analysis tool. AB, AG, PCTS and MC analyzed the data. PJC obtained funding and supervised TEVC experiments. NA designed, performed and analyzed TEVC experiments. AB and MC wrote the manuscript and all authors commented on it. All authors have read and agreed to the published version of the manuscript.

