## Supporting Information for "A millisecond coarse-grained simulation approach to decipher allosteric cannabinoid binding at the glycine receptor *α*1"

5 Predicted binding affinity of THC for GlyR using an increasing number of ligand copies in the simulation box at constant cannabinoid concentration. . 29

6 Total Number of Binding and Unbinding Events . . . . . . . . . . . . . . . . . 31

### Supplementary Method 1: Setup of an elastic network on the protein

The molecular system formed by the receptor, the lipid membrane, water molecules, counterions and NaCl 0.150 M (Supplementary Fig. 1) was simulated with an elastic network on the protein backbone to preserve its quaternary structure. The elastic network was introduced upon removing the endogenous neurotransmitter glycine from the orthosteric site. To search for an optimal strength of the harmonic restraint imposed by the elastic network (distance cutoff of 0.8 nm), multiple simulation runs with force constants ranging from 500 to 2500 kJ $mol^{-1}$ $nm^{-2}$ were carried out for 10 $\mu$s. Analysis of the backbone RMSD from the initial cryo-EM coordinates shows large deviations at low force constants, which plateau at $\sim$ 2 Å at significantly higher values (Supplementary Table 1). To preserve the receptor flexibility in the transmembrane domain, which is critical for ligand binding, the smallest value of $k_{elastic}$ that was able to maintain a symmetric architecture of the protein was chosen, i.e. $k_{elastic}$ of 1500 kJ $mol^{-1}$ $nm^{-2}$.

Supplementary Table 1: GlyR RMSD values with respect to the initial structure mapped to a CG representation by `martinize 2.0` as a function of the force constant, $k_{elastic}$, of the elastic network.

| **$k_{elastic}$ (kJ $mol^{-1}$ $nm^{-2}$)** | **RMSD (Å)** |
|---|---|
| 500 | 3.32 ± 0.19 |
| 750 | 3.31 ± 0.17 |
| 1000 | 2.68 ± 0.24 |
| 1250 | 2.41 ± 0.13 |
| 1500 | 2.39 ± 0.15 |
| 1750 | 2.20 ± 0.11 |
| 2000 | 2.11 ± 0.10 |
| 2250 | 2.03 ± 0.10 |
| 2500 | 2.08 ± 0.12 |

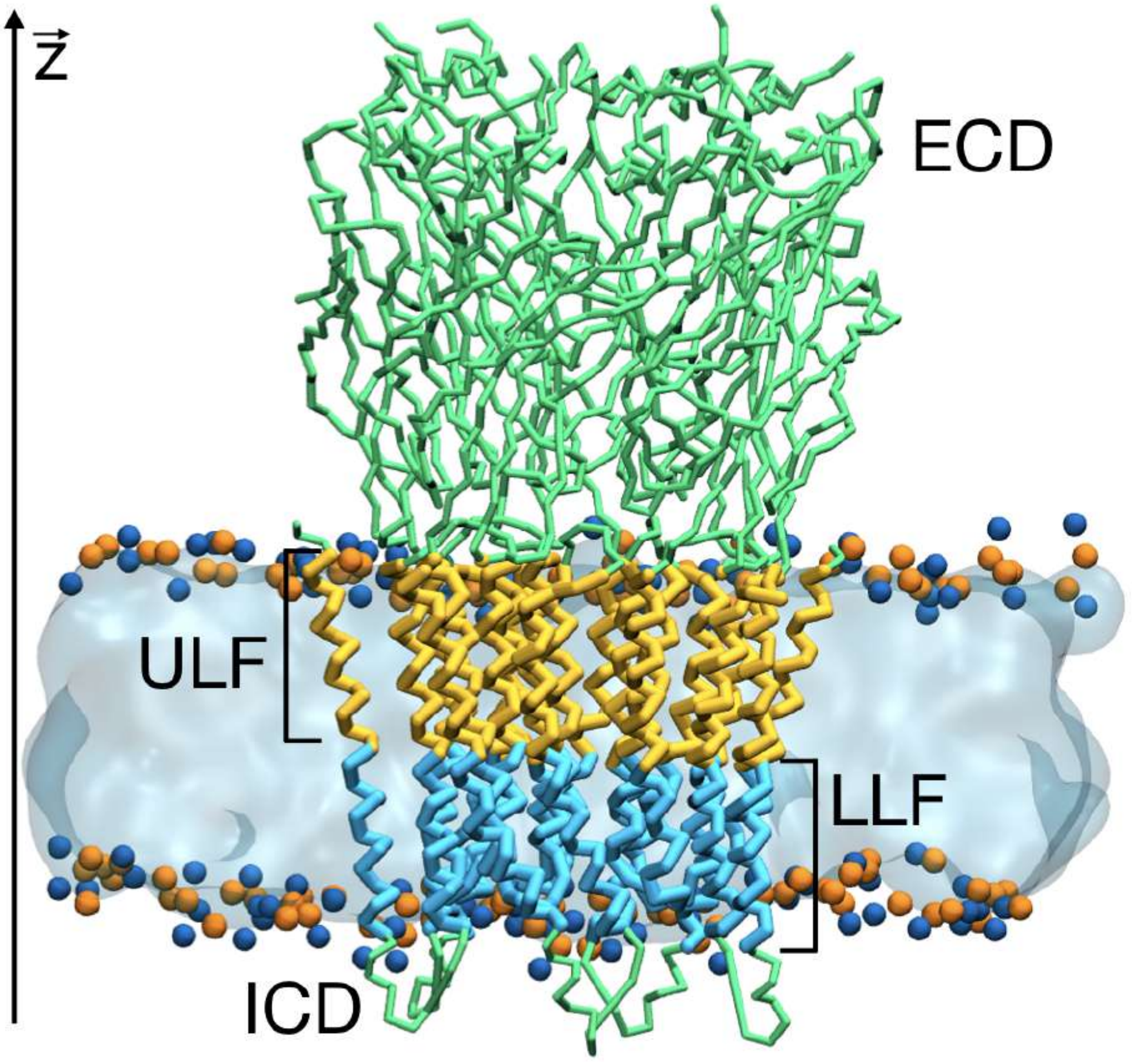

Supplementary Figure 1: Protein extracellular (ECD) and intracellular (ICD) domains are reported as green tubes. Membrane upper (ULF) and lower (ULF) leaflets are defined as the membrane regions along the Z axis perpendicular to the membrane, between the centers of mass of the whole POPC membrane (transparent cyan surface) and the POPC heads (NC3 and PO4 beads, reported as blue and orange spheres). The protein sections, comprised within the ULF and LLF membrane regions, are reported as orange and cyan tubes, respectively. Water and ions are not shown for the sake of clarity.

### Supplementary Method 2: Coarse–Grained Ligand Parameterization

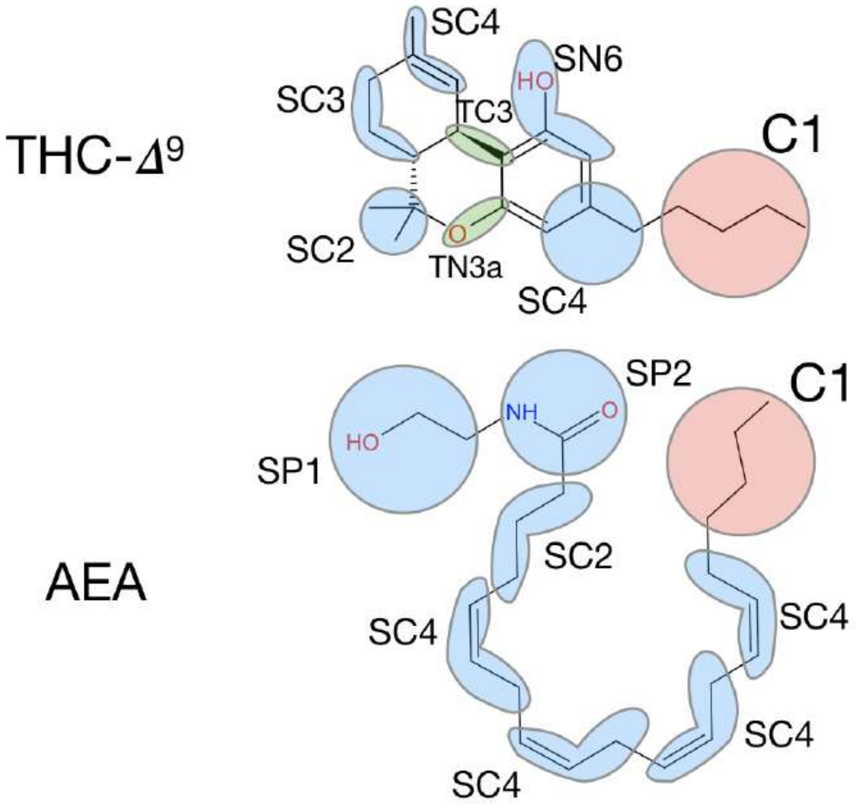

Supplementary Figure 2: THC and AEA coarse-grained mapping over the atomistic structure. Regular, small and tiny CG beads are colored in red, blue and green, respectively.

Ligand parameterization in Martini 3 involves several steps, as described in the work of Souza and co-workers.[1] In the following, we provide a summary of the procedure along with results obtained for THC and AEA.

#### Coarse-Grained mapping

The first step of the CG paramerization consists in mapping the atomic structure of the ligand into a set of interconnected beads. In this step, groups of atoms are selected to be represented by beads of different size (i.e. regular, small, and tiny). The CG mapping for THC and AEA used in this work is illustrated in Supplementary Fig. 2. This mapping relies on the combination of 4-1 (regular), 3-1 (small) and 2-1 (tiny) beads, with the goal of preserving chemical details and reproduce the overall molecular volume and shape as accurately as possible; see below.

#### Bonded Parameters

In the second step of the parameterization, all-atom MD simulations of the ligand in solution are carried out and the trajectories used to obtain reference distributions for all bead distances, angles and dihedrals based on the chosen CG mapping. By comparing with distributions obtained from CG/MD simulations, the force constants and equilibrium values for the bonded energy terms in the CG model are optimized to reproduce the all-atom results. Given that optimization of the bead distances and angles does not ensure an accurate description of the molecular shape and volume, an additional parameterization step was carried out where the solvent accessible surface area (SASA) and the Connolly surface of the CG model are compared with the all-atom MD results. In addition, structural analyses in solution based on the end-to-end distance and the radial distribution functions were performed as extra validation steps; see below.

##### All-Atom MD Simulations

Initial coordinates for THC and AEA were taken from the GRALL database[2] and converted into CHARMM36 topologies. The two cannabinoids were solvated with 4000 TIP3P water molecules, energy minimized for 50000 steps (steepest descent), equilibrated for 100 ns at 298 K (V-rescale thermostat, $\tau_T$ coupling constant of 0.1 ps) and 1 bar pressure (Berendsen barostat, $\tau_P = 5.0\,ps$, compressibility of $4.5 \cdot 10^{-5}\,bar^{-1}$), and simulated for 1.05 $\mu$s (of which the first 50 ns were discarded for analysis) using a Parrinello-Rahman barostat. Electrostatic interactions were treated using the PME method[3,4] (grid spacing of 0.10 nm and a PME-order of 4) with a cutoff of 1.2 nm, while the van der Waals interactions were cutoff at 1.2 nm. All bonds with hydrogens were constrained with the LINCS algorithm, which allows for use of an integration time step of 2 fs.

##### CG/MD Simulations

CG models of THC and AEA were solvated with 882 CG water beads, energy minimized and equilibrated for 20 ns at 298 K (V-rescale thermostat, $\tau_T$ coupling constant of 0.1 ps) and 1 bar (Berendsen barostat, $\tau_P = 4\,ps$, compressibility of $3 \cdot 10^{-4}\,bar^{-1}$), and simulated for 100 ns using a Parrinello-Rahman barostat. A 20 fs integration time step $t_{step}$ was used with periodic boundary conditions. Electrostatic interactions were cutoff at 1.1 nm using the reaction-field method, while Lennard-Jones interactions were cutoff at 1.1 nm using the potential-shift Verlet method. Bonds were constrained to their equilibrium values by using the LINCS algorithm.[5]

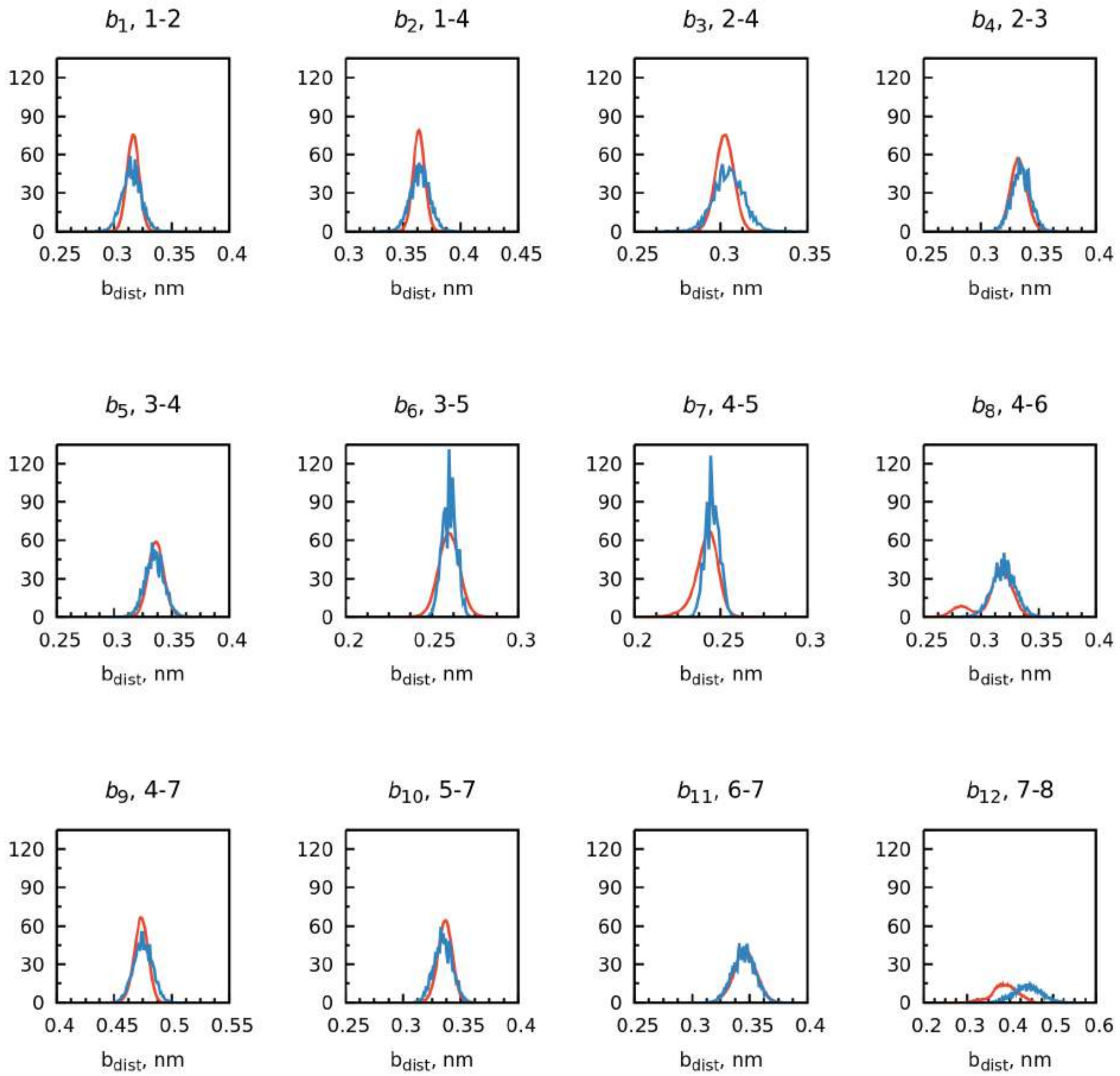

Supplementary Figure 3: THC distributions of bonds obtained from CG (blue line) and AA simulations (red line). In the AA simulation has been mapped the center-of-geometry (COG). In each plot title, *i*-*j* refers to the beads involved in such a bond term.

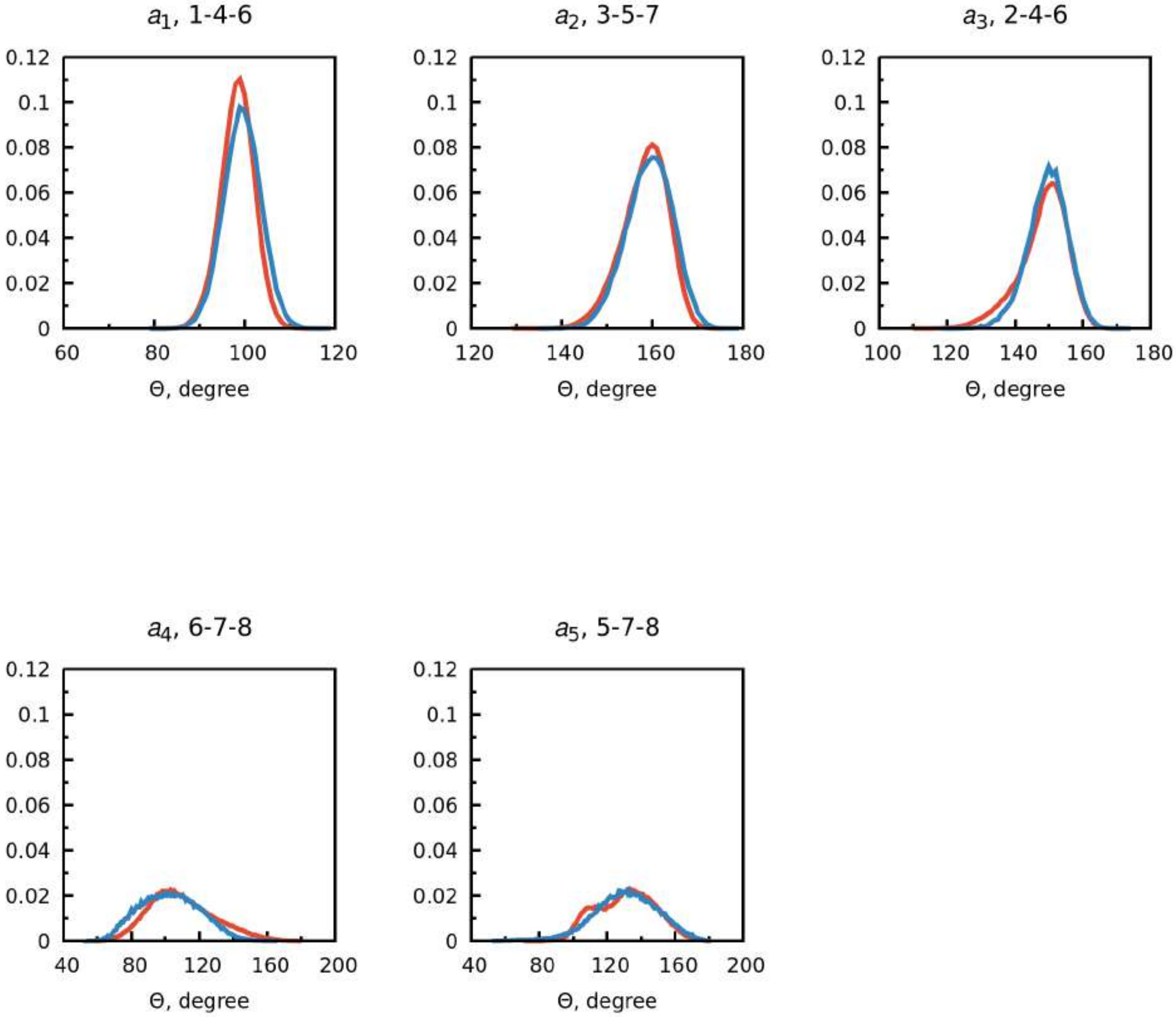

Supplementary Figure 4: THC distributions of angles obtained from CG (blue line) and AA simulations (red line). In the AA simulation has been mapped the center-of-geometry (COG). In each plot title, $i$-$j$-$k$ refers to the beads involved in such an angle term.

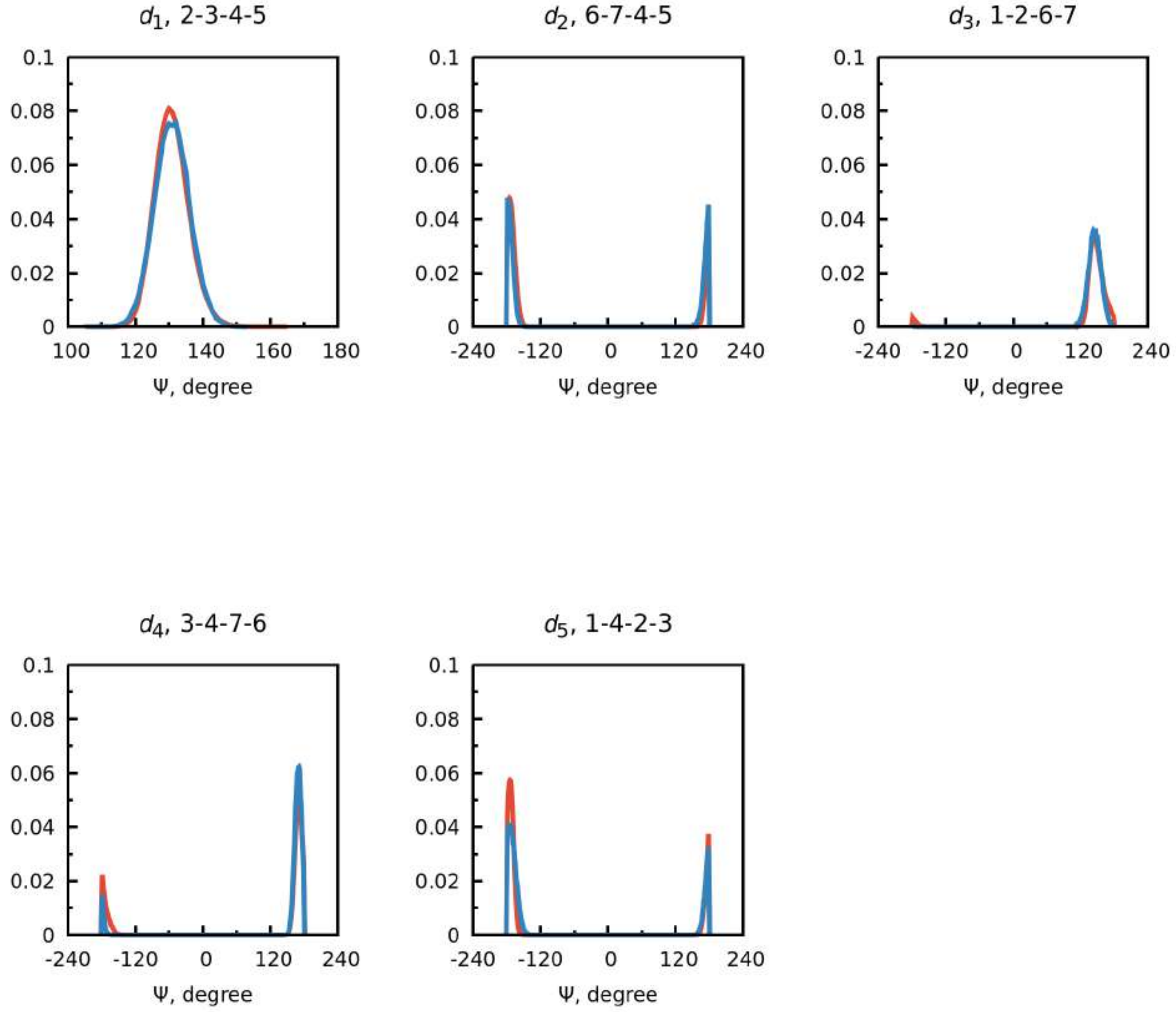

Supplementary Figure 5: THC distributions of dihedrals obtained from CG (blue line) and AA simulations (red line). In the AA simulation has been mapped the center-of-geometry (COG). In each plot title, $i$-$j$-$k$-$l$ refers to the beads involved in such a dihedral term.

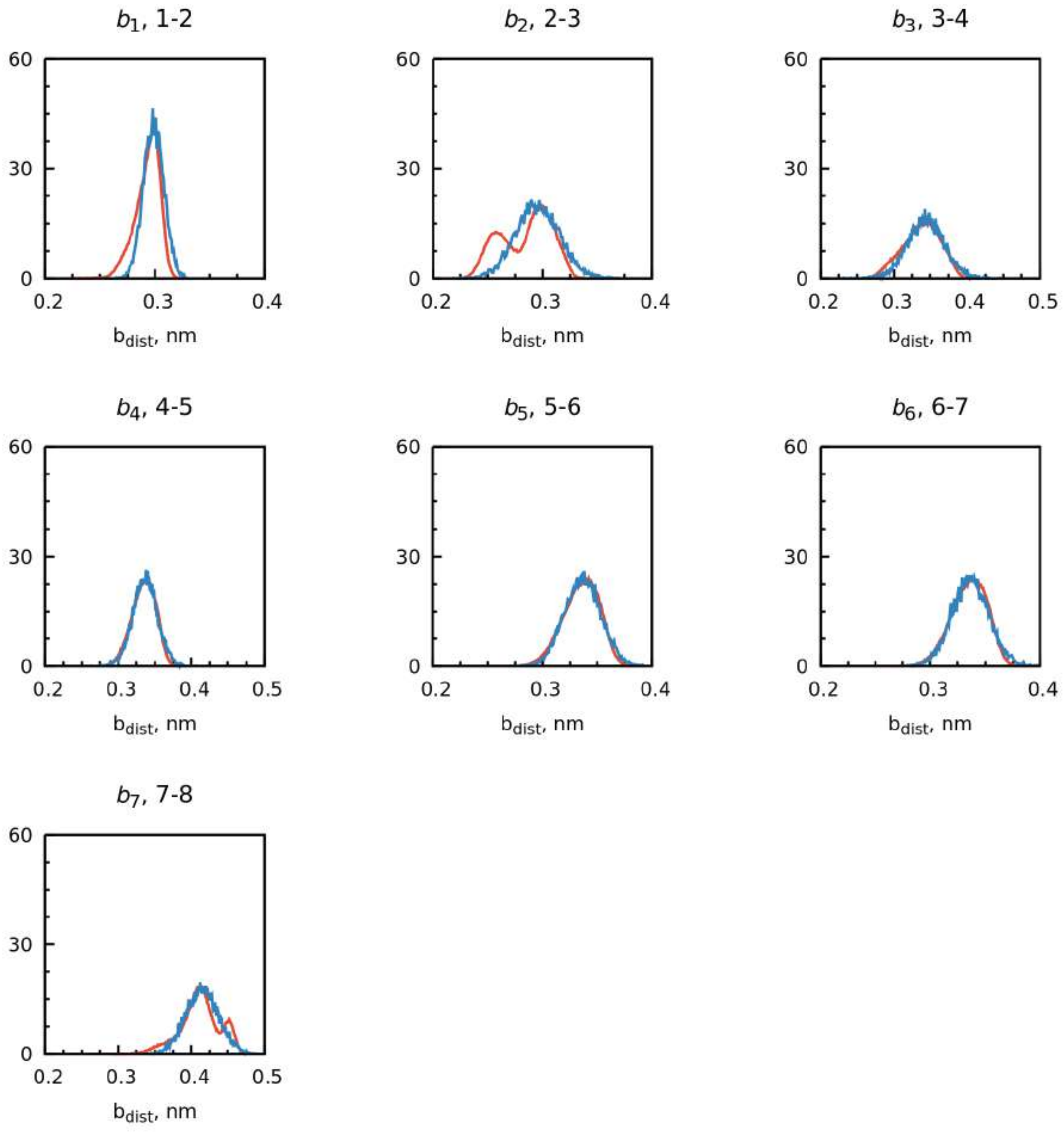

Supplementary Figure 6: AEA distributions of bonds obtained from CG (blue line) and AA simulations (red line). In the AA simulation has been mapped the center-of-geometry (COG). In each plot title, $i$-$j$ refers to the beads involved in such a bond term.

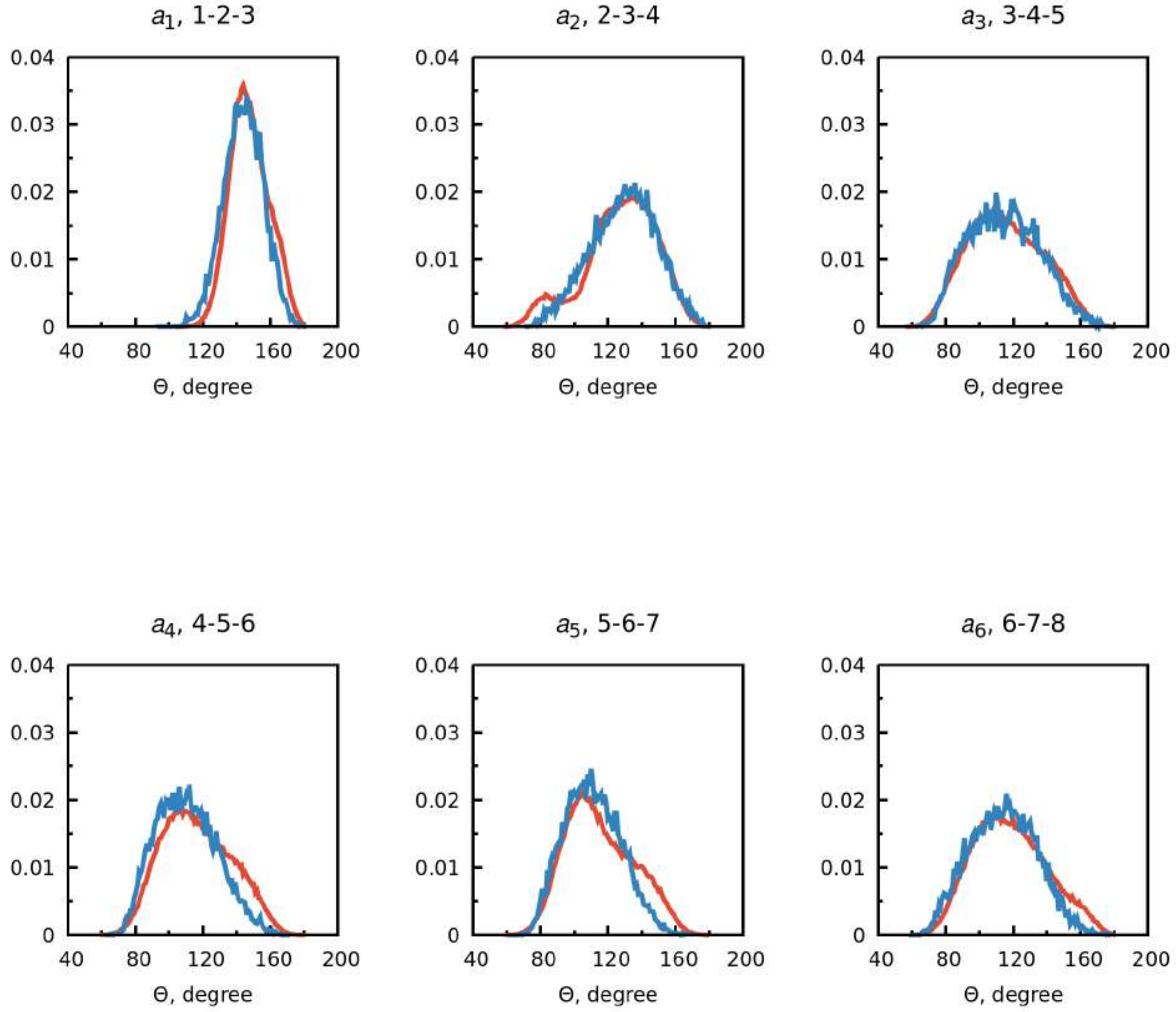

Supplementary Figure 7: AEA distributions of angles obtained from CG (blue line) and AA simulations (red line). In the AA simulation has been mapped the center-of-geometry (COG). In each plot title, $i$-$j$-$k$ refers to the beads involved in such an angle term.

#### Volume and Shape Refinement

The distribution of the SASA and the Connolly surface of THC and AEA from both all-atom MD and CG/MD simulations were obtained using the `gmx sasa` tool with a 0.191 nm probe radius and updated CG bead vdW-radii (R = 0.264 nm, S = 0.230 nm and T = 0.191 nm). The comparison is shown in Supplementary Fig. 8.

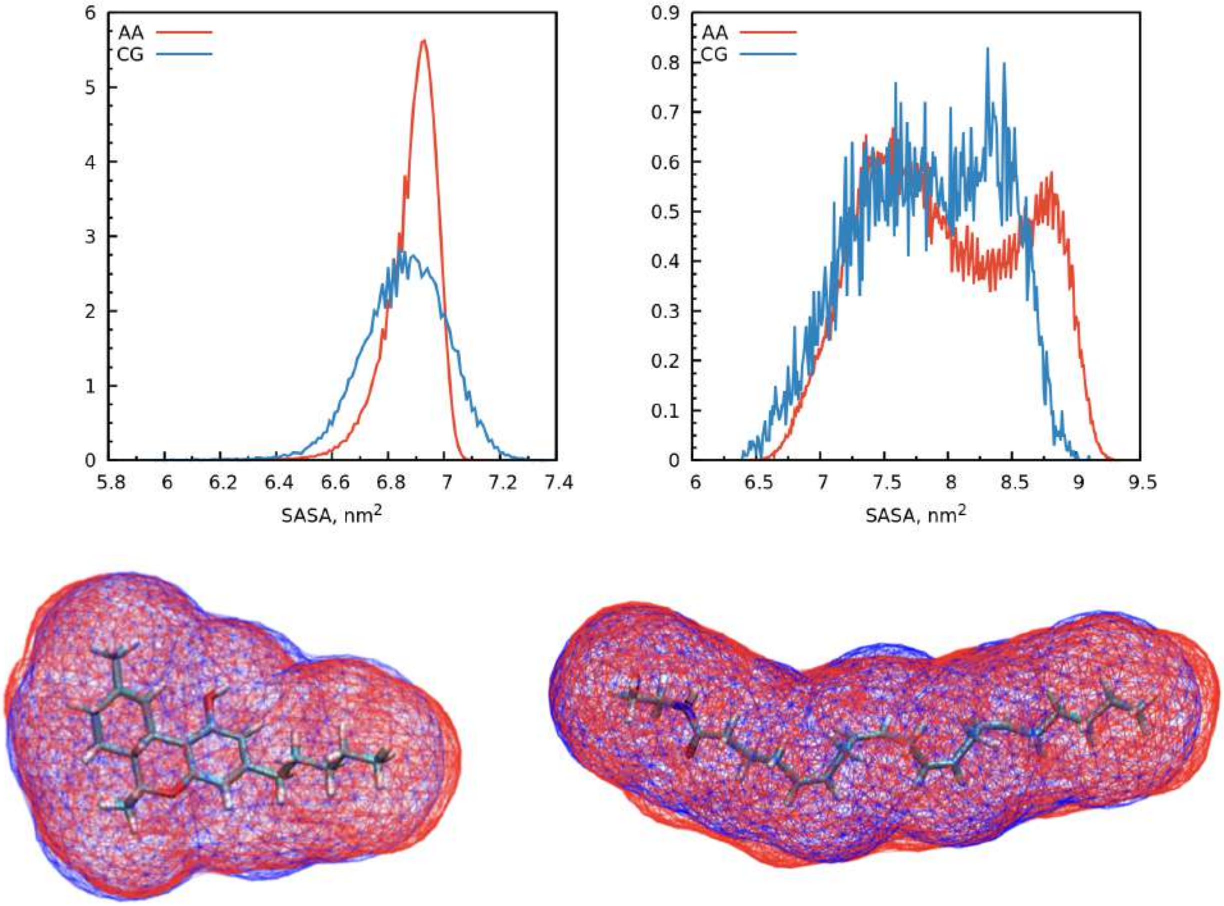

Supplementary Figure 8: **Upper panels:** distributions of SASA obtained from CG (blue line) and AA simulations (red line) for THC (left panel) and AEA (right panel). In the AA simulation has been mapped the center-of-geometry (COG). SASA average values are reported as it follows:
THC: 6.90 ± 0.10 $nm^2$ (AA) and 6.86 ± 0.15 $nm^2$ (CG);
AEA: 7.97 ± 0.61 $nm^2$ (AA) and 7.83 ± 0.55 $nm^2$ (CG)
**Lower panels:** Connolly surfaces for THC (left panel) and AEA (right panel) obtained from CG (blue wireframes) and AA simulations (red wireframes).

The molecular size of the ligand in solution was analyzed by comparing the end-to-end distance and the radial distribution functions (RDF, g(r)) using the `gmx gyrate` and `gmx rdf` tools. For the radial distribution function, the first bead (O1C in the AEA head and C1 in the THC cyclohexene ring) was chosen as reference point. The all-atom MD and CG/MD distributions are shown in Supplementary Figs. 9 and 10, respectively.

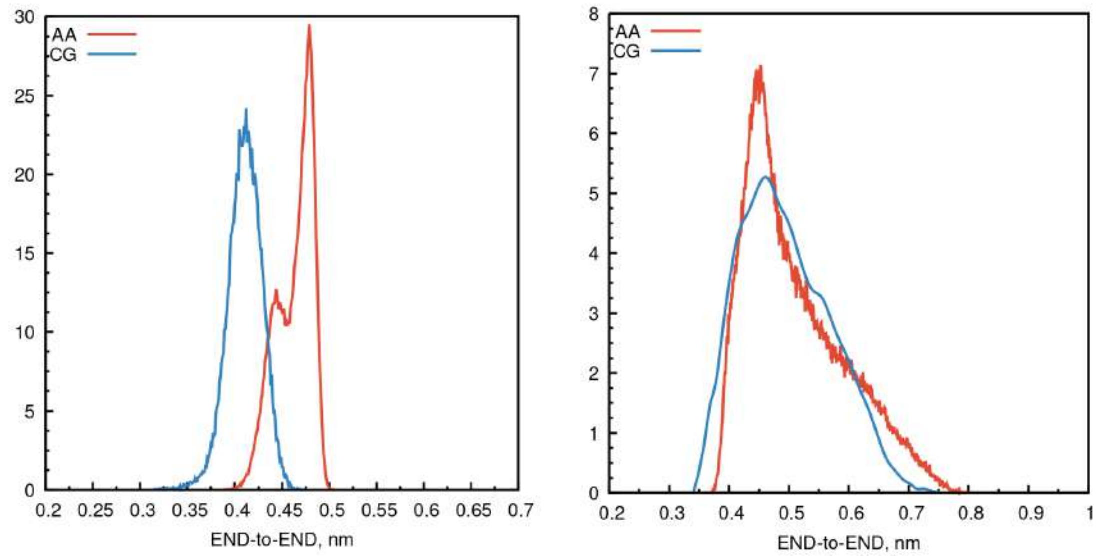

Supplementary Figure 9: Distributions of end-to-end distance from CG (blue line) and AA simulations (red line) for THC (left panel) and AEA (right panel). In the AA simulation has been mapped the center-of-geometry (COG). Average values of the end-to-end distance are reported as follows:
THC: 0.46 ± 0.02 nm (AA) and 0.41 ± 0.02 nm (CG)
AEA: 0.51 ± 0.08 nm (AA) and 0.50 ± 0.07 nm (CG)

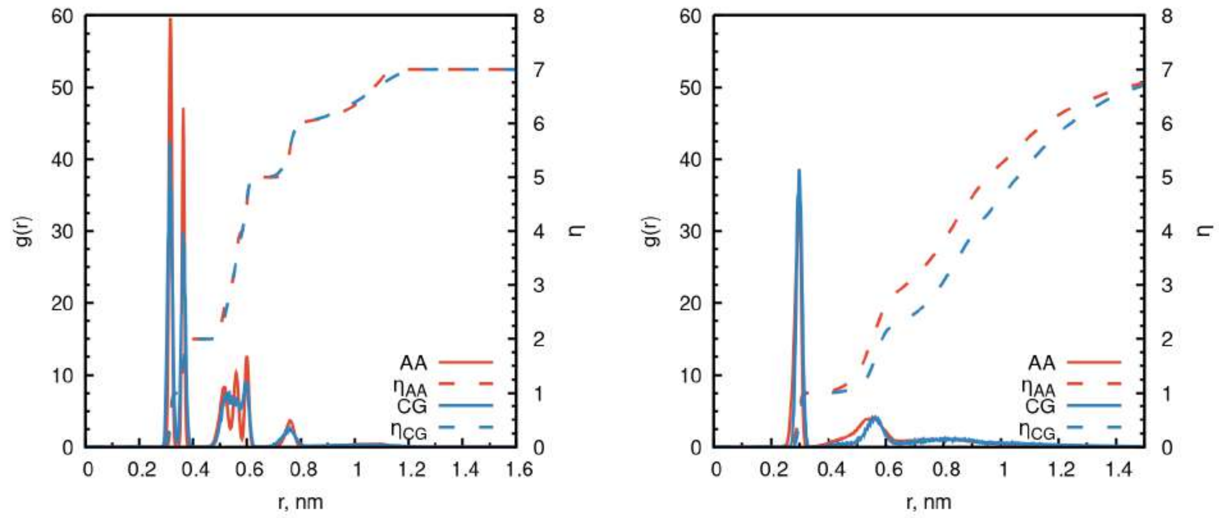

Supplementary Figure 10: Radial distribution functions (g(r)) from CG (blue line) and AA simulations (red line) for THC (left panel) and AEA (right panel). In the AA simulation has been mapped the center-of-geometry (COG). The dashed lines indicate the coordination number $\eta$, i.e. the average number of particles within a distance, r.

**Comparison of All-Atom MD with CG/MD Simulations**

By post-processing the production runs using the `gmx distance` and `gmx angles` tools and `gmx analyze`, the distributions of the bead distances, angles, and dihedrals in all-atom MD and CG/MD were obtained and compared. Equilibrium values and corresponding force constants for bonds, angles, and dihedrals were optimized iteratively by trials and errors until convergence to the all-atom MD results. The excellent agreement between the CG/MD (blue) and all-atom MD (red) distributions for both THC and AEA (Supplementary Figs. 3-7) validates the parameterization of the bonded interactions in the final CG models. In addition, the significant overlap between the distributions of the SASA (Supplementary Fig. 8), the end-to-end distance (Supplementary Fig. 9), and the radial distribution function (Supplementary Fig. 10) indicate that the CG models accurately reproduce the molecular surface and volume for the two cannabinoid derivatives.

#### Non-Bonded Parameters

The last step of the parameterization involves bead-type assignment, which controls the non-bonded interactions of the model. Initial bead types for THC and AEA were assigned based on the chemistry of the building blocks and taking inspiration from the extensive lits of examples provided in Table 1.1 of Alessandri et al.[6] and the Supplementary Tables 24 and 25 of Souza et al.[7] The parameterization was validated by computing the water/octanol partition coefficient ($\log P_{oct/wat}$) of the models and comparing it with literature data. The calculated $\log P_{oct/wat}$ for the CG models of THC and AEA was accessed from solvation free energy calculations, as follows. Starting from the water/octanol partitioning equilibrium

$$L_{wat} \rightleftarrows L_{oct}, \qquad \text{(Eq. S1)}$$

the definition of the partition coefficient

$$\log P_{oct/wat} = \log \left( \frac{[L]_{oct}}{[L]_{wat}} \right)_{eq}, \qquad \text{(Eq. S2)}$$

and the thermodynamic relation $\Delta G^\circ = -RT \ln K_{eq}$, it yields

$$\log P_{oct/wat} = -\frac{\Delta G^\circ_{wat \to oct}}{2.302\ RT} \qquad \text{(Eq. S3)}$$

Introducing the ligand in vacuum as an intermediate for the water to octanol transfer reaction, Eq. S3 can be rearranged as

$$\log P_{oct/wat} = -\frac{\Delta G^\circ_{wat \to vac} - \Delta G^\circ_{oct \to vac}}{2.302\ RT} \qquad \text{(Eq. S4)}$$

which indicates that $\log P_{oct/wat}$ can be predicted from the desolvation free energy of the ligand in water and in octanol, which are quantified by alchemical free energy calculations; see below. And, using Eq. S4, the uncertainty on the calculated $\log P_{oct/wat}$ can be estimated as

$$\delta \log P_{oct/wat} = \frac{\sqrt{\delta \Delta G^{\circ}_{wat \to vac}{}^2 + \delta \Delta G^{\circ}_{oct \to vac}{}^2}}{2.302\ RT} \qquad \text{(Eq. S5)}$$

#### Solvation Free Energies of CG Models

To evaluate their solvation free-energy, CG models of THC or AEA were solvated with 882 water beads in water solution, or 869 octanol and 54 water beads in a 25 % octanol-water mixture[8]) as prescribed by standard protocols in Martini 3.[6,7,9] The solvation free energy of the ligand was computed by thermodynamic integration (TI) using 23 $\lambda$-windows to decouple the solute-solvent vdW interactions following the $\lambda$-schedule: $\lambda_{vdW}$=(0.000, 0.050, 0.100, 0.200, 0.300, 0.400, 0.500, 0.600, 0.625, 0.650, 0.675, 0.700, 0.725, 0.750, 0.775, 0.800, 0.825, 0.850, 0.875, 0.900, 0.925, 0.950, 1.000). After an initial minimization of 50000 steps (steepest descent), the simulations were carried out for 60 ns per window and $\left(\frac{\partial H}{\partial \lambda}\right)_{NpT,\lambda}$ were collected after 10 ns equilibration (i.e. 50 ns per window). In the implemented approach, the ligand interacts with the solvent in the initial state ($\lambda = 0$), while it is fully decoupled (i.e. isolated in vacuum) in the final state ($\lambda = 1$). To avoid numerical problem, a soft-core potential[10] with $\alpha = 0.5$, power = 1 and $\sigma = 0.3$ was used. For all free energy simulations, 20 fs integration time step with periodic boundary conditions and the `sd` (i.e. stochastic dynamics) integrator in gromacs were used. During the equilibration and production runs, the temperature was maintained constant at 298 K with a V-rescale thermostat[11] with a $\tau_T$ coupling constant of 1 ps. The pressure was kept constant at 1 bar (isotropic coupling) with a Parrinello-Rahman barostat[12] ($\tau_P = 12\,ps$, compressibility of $3 \cdot 10^{-4}\,bar^{-1}$). To quantify the solvation free energy and associated error, the Multistate Bennett Acceptance

Ratio (MBAR) was used as implemented in `alchemical-analysis.py`.[13] The free-energy results in water and octanol for THC and AEA are given in Supplementary Table 2. The calculated $\log P$ for THC reproduces the experimental determination[14,15] within one log unit. The $\log P$ for AEA deviates some more from the literature, which is based on predictions from machine learning,[16] but the trend is consistent; i.e., the CG model for AEA is less hydrophobic than that for THC.

Supplementary Table 2: THC and AEA desolvation free–energies (in kcal/mol) and water/octanol partition coefficient $\log P_{oct/wat}$ at 298 K at the CG level.

| **System** | $\Delta G^{\circ}_{wat \rightarrow vac}$ | $\Delta G^{\circ}_{oct \rightarrow vac}$ | $\log P^{calc}_{oct/wat}$ | $\log P^{ref}_{oct/wat}$ |
|---|---|---|---|---|
| THC | $1.000 \pm 0.016$ | $11.836 \pm 0.015$ | $7.950 \pm 0.016$ | 6.97 (HPLC[14,15]) |
| AEA | $5.030 \pm 0.018$ | $14.549 \pm 0.016$ | $6.980 \pm 0.018$ | 5.1 (ref.[16]) |

### Supplementary Method 3: Protein-Ligand Binding at Equilibrium

In this section, we derive an expression that is straightforward to use for the evaluation of the standard free energy of binding from equilibrium CG/MD simulations. First, we present the theory and derive the expression. Then, we describe the implementation that was used to estimate the binding affinity of THC and AEA for GlyR in *Main Text*. Last, we present model calculations for THC in interaction with GlyR to show that this expression is independent of the ligand concentration or the number of ligands in the simulation box. Remarkably, the results show that CG/MD simulations may provide binding affinity predictions with a statistical uncertainty $< 0.1$ kcal/mol in hundreds of $\mu$s sampling. We conclude that CG/MD and Eq. S15 provide straightforward access to the ligand $K_d$ even for complex molecular systems involving transmembrane proteins embedded in lipid bi-layers.

#### Theory

Consider the following chemical reaction

$$P + L \rightarrow PL \qquad \text{(Eq. S6)}$$

with $P$ being a monomeric or heteromeric protein molecule, $L$ a small-molecule ligand, and $PL$ the protein-ligand complex formed by non-covalent interactions in solution. The difference in chemical potential between reactants and products is

$$\Delta\mu_{bind} = \mu_{PL} - \mu_P - \mu_L \qquad \text{(Eq. S7)}$$

In the limit of idealized solution behaviour (i.e., particles are independent and indistinguishable), the chemical potential of each reaction component can be expressed as

$$\mu_i = -kT \ln \frac{q_i}{N_i} \qquad \text{(Eq. S8)}$$

with $q_i$ and $N_i$ being the molecular partition function and the number of molecules of the $i$-th species. Introducing this result into Eq. S7, it yields

$$\Delta\mu_{bind} = -kT \ln \left( \frac{q_{PL}}{q_P \times q_L} \right) + kT \ln \left( \frac{N_{PL}}{N_P \times N_L} \right) \qquad \text{(Eq. S9)}$$

At chemical equilibrium, $\Delta\mu_{bind} = 0$ and Eq. S9 yields[17]

$$\left( \frac{N_{PL}}{N_P \times N_L} \right)_{eq} = \left( \frac{q_{PL}}{q_P \times q_L} \right) \qquad \text{(Eq. S10)}$$

By dividing both sides of the equation by the solution volume, $V$, and the Avogadro constant, $N_A$, it yields

$$\left( \frac{C_{PL}}{C_P \times C_L} \right)_{eq} = \frac{q_{PL}/V}{(q_P/V)(q_L/V)} N_A \qquad \text{(Eq. S11)}$$

with $C_i$ being molar concentrations, i.e. moles per liter. Because in the ideal gas approximation molecular partition functions are of the form $f(T) \times V$, the right-hand side of Eq. S11 is volume independent and is a chemical equilibrium constant

$$K_d^{-1} = \frac{q_{PL}}{q_P \times q_L} V N_A \qquad \text{(Eq. S12)}$$

The result of Eq. S12 shows that the binding constant has units of L/mol. Assuming that the ligand is small and there is no conformational change of the protein upon binding, the binding reaction can be understood as a ligand partitioning equilibrium ($L_u \rightleftharpoons L_b$ ), and the partition function ratio in Eq. S12 be approximated by $\frac{q_{L,b}}{q_{L,u}}$, with $q_{L,b}$ and $q_{L,u}$ being the molecular partition function of the ligand in the bound and unbound states, respectively. At chemical equilibrium (Eq. S10)

$$\left(\frac{N_{L,b}}{N_{L,u}}\right)_{eq} = \frac{q_{L,b}}{q_{L,u}} = \frac{p_b}{p_u} \tag{Eq. S13}$$

and the partition function ratio can be measured from the equilibrium probabilities for the ligand to be bound ($p_b$) or unbound ($p_u$). Note that since $q_{L,u}$ is a function of volume, the probability of binding is volume (or concentration) dependent and approaches zero at infinite dilution. Introducing this result in Eq. S12, it yields

$$K_d^{-1} = \frac{p_b}{p_u} V N_A \tag{Eq. S14}$$

which has units of L/mol. By introducing an arbitrary standard concentration $C^\circ = n^\circ / V^\circ$, i.e. typically 1 mol/L, the product $K_d^{-1} C^\circ$ is made dimensionless and the standard free energy of binding is obtained as

$$\Delta G^\circ_{\text{bind}} = -RT \ln \left( \frac{p_b}{p_u} \frac{n^\circ N_A V}{V^\circ} \right) \tag{Eq. S15}$$

As shown by Eq. S15, the numerical value of $\Delta G^\circ_{\text{bind}}$ depends on the definition of the standard state, which sets the number of molecules ($N^\circ = n^\circ N_A$) and the solution volume ($V^\circ$) in this reference state, as well as the units of measure for the solution volume in the actual experiment ($V$). We note that the result of Eq. S15 is rigorous only in the absence of molecular processes other than protein-ligand binding such as ligand or protein aggregation. We also note that the same expression was derived for dimerization reactions based on slightly different theoretical considerations.[18]

#### Implementation

The result of Eq. S15 indicates that protein-ligand binding affinities ($\Delta G^\circ_{\text{bind}}$) can be directly accessed from the probability of the ligand to be bound ($p_b$) and the solution volume ($V$). As illustrated below, all these quantities can be extracted from a converged MD simulation, which thus provides straightforward access to the binding energetics. This is particularly true in the simulated conditions used in this work, where only one receptor is modeled (no protein aggregation) and ligand aggregation is only marginally observed as hydrophobic molecules such as THC and AEA tend to be dispersed in the lipid membrane.

##### Probability of binding

In our implementation of Eq. S15, the probability for the ligand to be bound ($p_b$) or unbound ($p_u$) were estimated using a contact analysis of the CG/MD trajectories. The time series of the protein-ligand contacts was generated using the `gmx mindist` tool with a cutoff distance of 0.6 nm. All simulation frames with the number of contacts larger than unity were considered as representatives of the bound state, whereas those with zero contacts as representatives of the unbound state. The values of $p_b$ and $p_u$ were estimated by collecting statistics over multiple ligands in the simulation box and several simulation replicas.

##### Quantification of the volume

The solution volume ($V$) in Eq. S15 is the volume that is accessible to the ligand. For a monophasic system like a soluble protein, $V$ simply corresponds to the volume of the simulation box minus the volume of the protein. For a biphasic system like a transmembrane protein embedded in a lipid bilayer, the effectively accessible volume is impacted by ligand partitioning among the two phases and becomes closer to the membrane volume for highly hydrophobic compounds as the cannabinoids. In this work, the solution volume was estimated from the average size of the simulation box along the X, Y, and Z axes minus the volume of the protein as quantified by the Voss Volume Voxelator software[19] (`vossvolvox`, version 1.2). Using a probe radius of 3Å and the grid quality set to "High", the volume of the GlyR active state (6PM6) was estimated to be 295.9 nm$^3$. Values of the solution volume ($V$) and the molar concentration ($C$) of cannabinoids in the CG/MD simulations are given in Supplementary Table 3. To estimate the increase in the effective concentration

Supplementary Table 3: Volume and ligand concentration in the CG/MD simulations of THC and AEA in interaction with GlyR.

| System | N | X (nm) | Y (nm) | Z (nm) | $V_{box}$ (nm$^3$) | V (nm$^3$) | C (M) | $V_{memb}$ (nm$^3$) | $C_{memb}$ (M) |
|---|---|---|---|---|---|---|---|---|---|
| THC | 16 | 11.534 | 11.534 | 19.721 | 2624 | 2328 | 0.011 | 397 | 0.067 |
| AEA | 16 | 11.571 | 11.571 | 19.580 | 2622 | 2326 | 0.011 | 400 | 0.066 |

of cannabinoid by ligand partitioning in the membrane, the effectively accessible volume was quantified as the volume of the membrane bilayer ($V_{memb}$) minus the volume of the transmembrane domain of the protein ($V_{TMD}$). The membrane volume was determined as

$$V_{memb} = A_{bilayer} \times d_{bilayer} \qquad \text{(Eq. S16)}$$

where $A_{bilayer}$ was taken as the product of the box sizes parallel to the membrane plane (XY), and $d_{bilayer}$ was estimated from the statistical distribution of the lipid heads in the direction perpendicular to the membrane (Z). For this purpose, the distribution of the lipid heads was computed using the `gmx density` tool over 10 $\mu$s of CG/MD simulation, and $d_{bilayer}$ quantified from the distance between the two main density peaks; e.g., in our pure POPC bilayer this distance was 3.85 nm. The volume of the TMD of the protein was determined by `vossvolvox` using a PDB structure containing all protein atoms whose Z coordinate was between the upper and lower leaflet boundaries; see Supplementary Fig. 11. The volume of the TMD in the active state of GlyR was estimated to be 115.1 nm$^3$, which is approximately one third of the protein volume. Using $V_{memb}$ as the ligand-accessible volume, the effective concentration of cannabinoids in our experiments increases six-fold; see Supplementary Table 3.

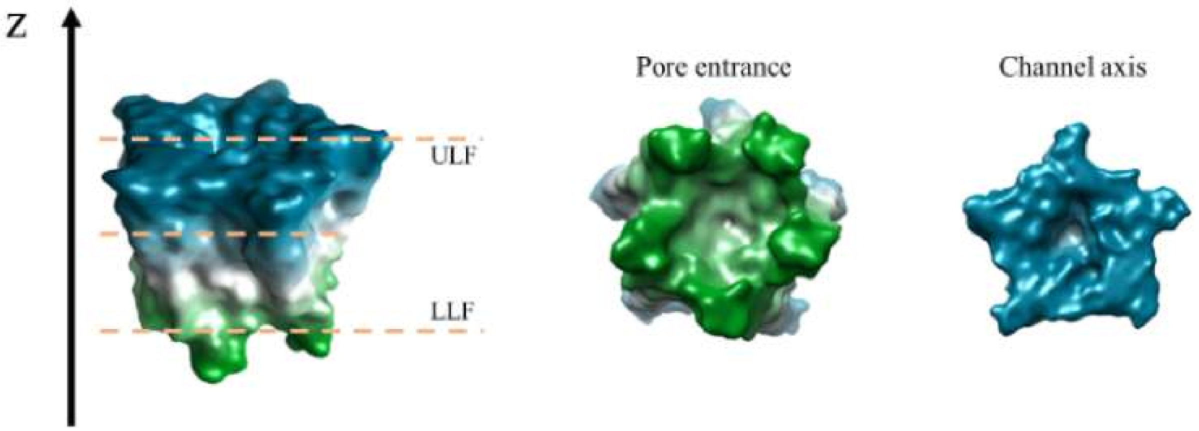

Supplementary Figure 11: Example of the 3V volume analysis applied to an atomistic .pdb file of the transmembrane domain of the Glycine receptor. Dashed lines represent upper and lower boundaries of the lipidic bilayer (upper leaflet, ULF; lower leaflet, LLF).

#### Error estimate

Eq. S15 states that the protein-ligand binding affinity can be accessed from the probability of the ligand to be bound, which is quantified from converged CG/MD simulations. If so, the error on the binding affinity is solely due to the uncertainty on the binding probability ratio extracted from the simulations. Since the uncertainty on the binding probability ratio is $\frac{\delta(p_b)}{(1-p_b)^2}$ from error propagation [1] and that on $\ln x \pm \delta x \approx \frac{\delta x}{x}$, assuming that $\delta p_b$ can be estimated from the standard error of the mean ($S.E.M.$) computed over multiple simulation replicas, the uncertainty on the binding affinity is

$$\delta \Delta G^{\circ}_{\text{bind}} = RT \, \frac{S.E.M.}{p_b(1-p_b)} \qquad \text{(Eq. S17)}$$

[1] $\delta\left(\frac{p_b}{p_u}\right) = \left(\frac{\delta(p_b)}{p_b} + \frac{\delta(p_u)}{p_u}\right)\frac{p_b}{p_u} = \left(\frac{\delta(p_b)}{p_b} + \frac{\delta(p_b)}{1-p_b}\right)\frac{p_b}{1-p_b} = \frac{\delta(p_b)}{(1-p_b)^2}$

#### Application to GlyR-THC

##### Dependence on the ligand concentration

The influence of the initial concentration of THC on the predicted affinity for GlyR was explored by simulating one protein with one ligand in a set of boxes with increasing size; see Supplementary Fig. 12. For each system, 10-$\mu$s long CG/MD simulations were carried out

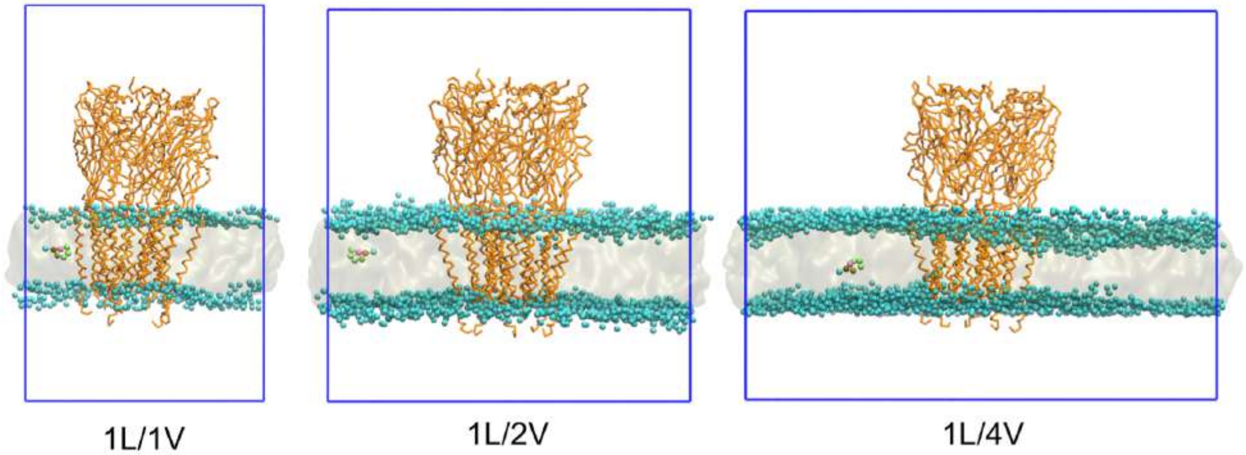

Supplementary Figure 12: Exploration of the THC-binding affinity for GlyR at different concentrations of cannabinoids. The simulation boxes used in these model calculations are shown. The protein is shown in orange, the lipid heads in light blue, and the THC ligand in colors. In these experiments, the volume of the simulation box used in *Main Text* (left) was increased $\times 2$ (middle) and $\times 4$ (right) along the XY axes.

in 20 replicas. The ligand-binding affinities predicted by Eq. S15 are given in Supplementary Table 4. The data demonstrate that the result of Eq. S15 is concentration independent and the statistical uncertainty based on 200 $\mu$s sampling is $< 0.1$ kcal/mol. Based on these calculations, we conclude that the affinity of our model THC for GlyR is $\sim 10$ mM.

Supplementary Table 4: Predicted binding affinity of THC for GlyR at decreasing cannabinoid concentrations but same number of ligand copies in the simulation box.

| Experiment | No. THC | V ($nm^3$) | C (M) | $p_b$ | $p_u$ | $\Delta G^{\circ}_{\text{bind}}$ (kcal/mol) | $K_d$ (mM) |
|---|---|---|---|---|---|---|---|
| 1L/1V | 1 | 2336 | 0.0007 | 0.23 | 0.77 | $-3.60 \pm 0.03$ | 2.4 |
| 1L/2V | 1 | 4493 | 0.0004 | 0.11 | 0.89 | $-3.46 \pm 0.07$ | 3.0 |
| 1L/4V | 1 | 8988 | 0.0002 | 0.07 | 0.93 | $-3.56 \pm 0.07$ | 2.5 |

**Dependence on the number of ligands**

To explore the influence of the number of simulated ligands on the binding affinity prediction by Eq. S15, a series of model systems with increasing simulation boxes but same nominal concentration of THC were explored; see Supplementary Fig. 13. For this purpose, unbiased CG/MD simulations were carried out for 10 $\mu$s for each molecular system and in 20 replicas. The results are given in Supplementary Table 5. The data demonstrate that the binding affinity predicted by Eq. S15 are independent of the number of simulated ligands, at least at sub-saturating concentrations of THC; i.e., when the number of simulated ligands is less than the number of binding sites on the protein. Given the close agreement between the binding affinity results in Supplementary Table 5 with those given in Table 1 of *Main Text* for THC, we conclude that 5% THC or AEA is not far from the idealised solution behaviour, which justifies the use of Eq. S15 for all binding affinity determinations.

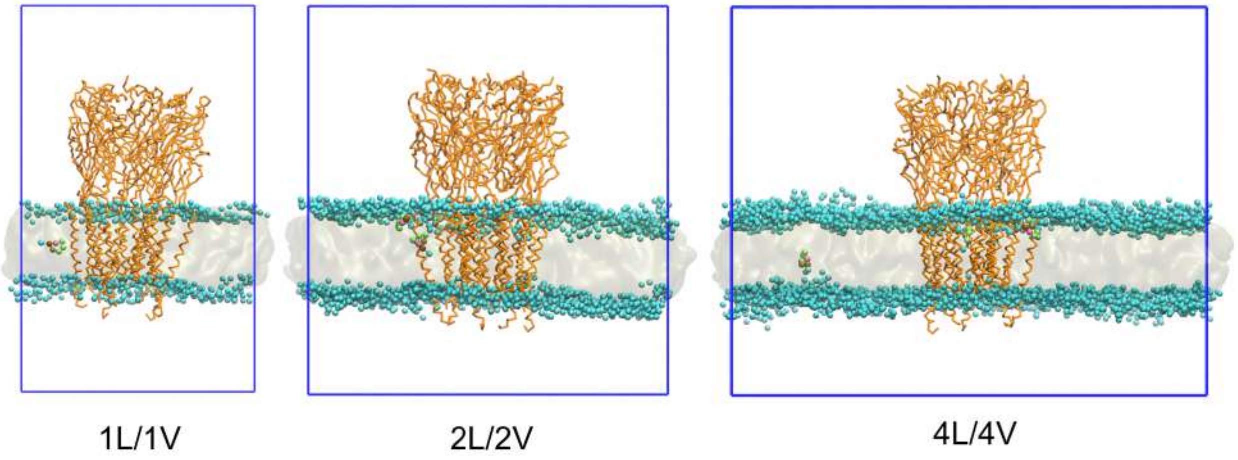

Supplementary Figure 13: Exploration of the THC-binding affinity for GlyR using an increasing number of ligand copies in the simulation box at constant cannabinoid concentration. The simulation boxes used in these model calculations are shown. To maintain the same cannabinoid concentration, the number of THC ligands was increased proportionally to the volume of the lipid bilayer. The color code is the same as in Supplementary Fig. 12

.

Supplementary Table 5: Predicted binding affinity of THC for GlyR using an increasing number of ligand copies in the simulation box at constant cannabinoid concentration.

| Experiment | No. THC | V ($nm^3$) | C (M) | $p_b$ | $p_u$ | $\Delta G^\circ_{\text{bind}}$ (kcal/mol) | $K_d$ (mM) |
|---|---|---|---|---|---|---|---|
| 1L/1V | 1 | 2336 | 0.0007 | 0.23 | 0.77 | $-3.60 \pm 0.03$ | 2.4 |
| 2L/2V | 2 | 4493 | 0.0007 | 0.12 | 0.88 | $-3.52 \pm 0.03$ | 2.7 |
| 4L/4V | 4 | 8692 | 0.0008 | 0.07 | 0.93 | $-3.60 \pm 0.05$ | 2.4 |

### Supplementary Method 4: GlyR-Ligand Interaction

#### Binding and Unbinding Events

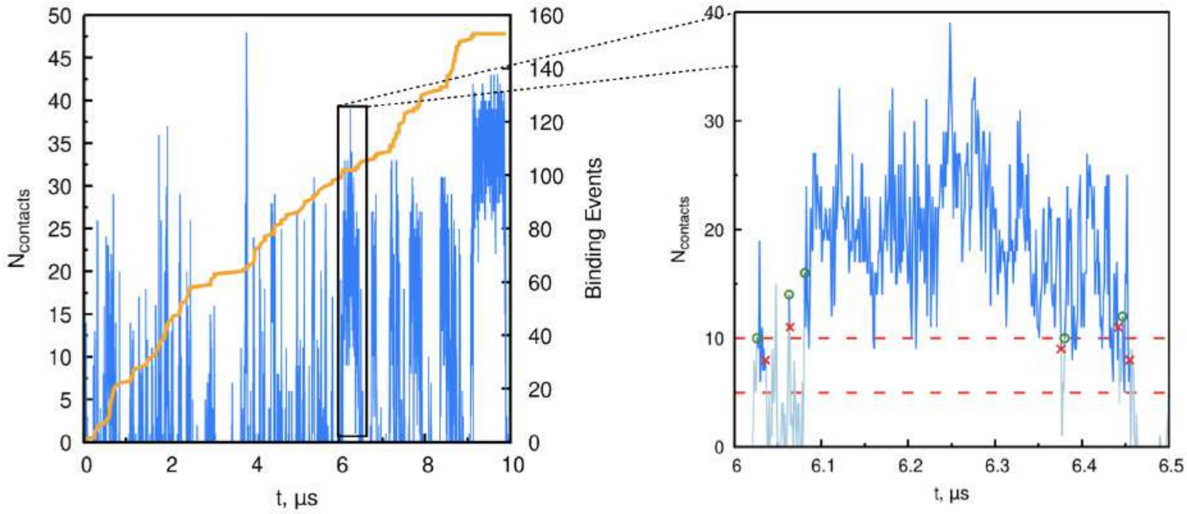

Supplementary Figure 14: On the left, the time series of the number of contacts between one AEA molecule and GlyR during 10 $\mu s$ simulation is shown in blue. The number of AEA-binding events is shown in orange. On the right, a zoom of 0.5 $\mu s$ simulation is used to illustrate the dual-cutoff scheme used to identify ligand-binding events. Both the upper (10 contacts) and lower (5 contacts) cutoffs are shown as red dashed lines. The ligand-binding event (green circle) starts when the number of contacts is $\geq$ 10 (i.e., the upper cutoff). When the number of contacts becomes $<$ 5 (i.e., the lower cutoff) the binding event stops (red cross). The dual-cutoff is used to handle the statistical noise on the number of ligand-receptor contacts.

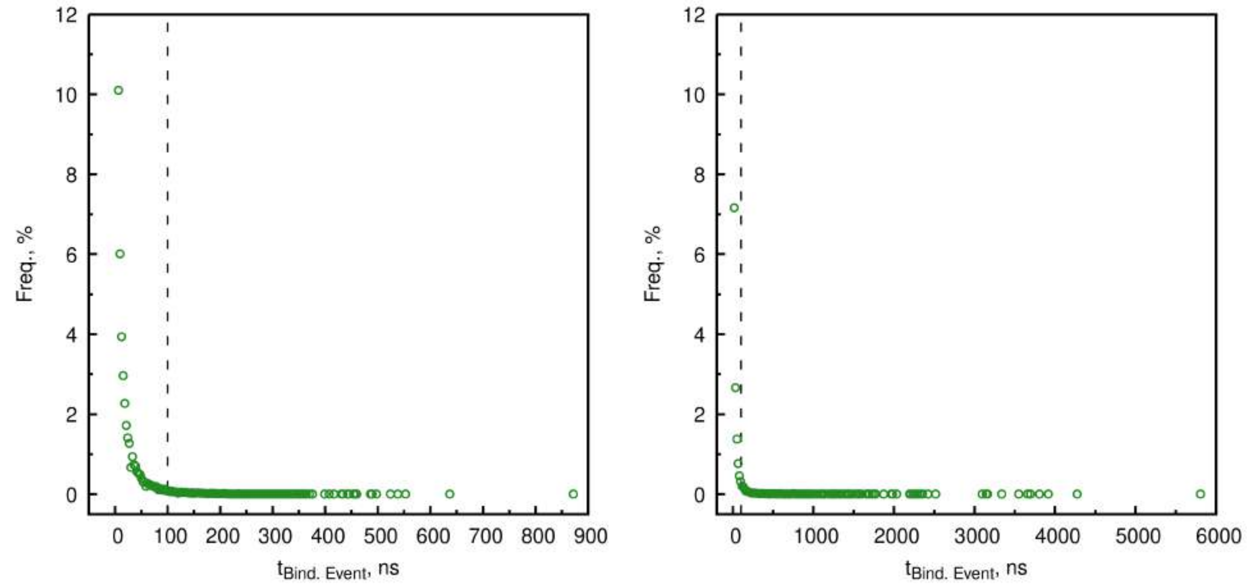

Supplementary Figure 15: Residence times ($t_{Bind.\,Event}$, in ns) distributions of GlyR-THC (left panel) and GlyR-AEA (right panel) binding events. The vertical dashed line at 100 ns indicates the cutoff for selecting specifc ($\tau_r \geq 100$ ns) and non-specific events ($\tau_r < 100$ ns), as described in Methods. The total number of binding (specific and non-specific) and unbinding events is reported in Supplementary Table 6. The number of bins in the exponential distribution is equal to $\sqrt{N_{bindingevents}}$.

Supplementary Table 6: Total number of binding (specific and non–specific) and unbinding events for GlyR-THC and GlyR-AEA systems.

| **System** | **Binding Events** | | | **Unbinding Events** |
|---|---|---|---|---|
| | **Total** | **Specific** | **Non–specific** | |
| GlyR-THC | 91266 | 1290 | 89976 | 91235 |
| GlyR-AEA | 129346 | 1880 | 127466 | 129283 |

#### Binding Modes

The decomposition of the CG/MD simulation trajectories into ligand-binding events allows to deconvolute the 2D and 3D maps shown in Fig. 1 of *Main Text* into contributions per binding mode. 2D-density maps were obtained using the `gmx densmap` tool, while 3D-density maps using the `volmap` plugin of `VMD`.[20] The radius of each ligand bead was set to 2.64 Å, 2.30 Å and 1.91 Å for the regular (R), small (S) and tiny (T) bead size. 2 Å was used as volume cube side length (`-res` option). The results for THC and AEA are shown in Supplementary Fig. 16 and Supplementary Fig. 17, respectively. For THC, the deconvolution of the 2D density shows that: i. the density peaks in Fig. 1 of *Main Text* well correspond to contributions from the three dominant binding modes (orange, red, and cyan in Fig. 2); and ii. intrasubunit binding corresponds to the brightest spot in Fig. 1 because both mode 2 (red) and mode 3 (cyan) contribute to it. For AEA, it shows that: i. the deeper penetration of the ligand in the upper leaflet in Fig. 1 of *Main Text* actually belongs to intrasubunit binding, in particular mode 1 (green); ii. the deeper penetration of the ligand in the lower leaflet in Fig. 1 corresponds to lower intersubunit binding (red); and iii. the broad density peaks in the upper leaflet in Fig. 1 originate from upper intersubunit binding.

#### THC Results

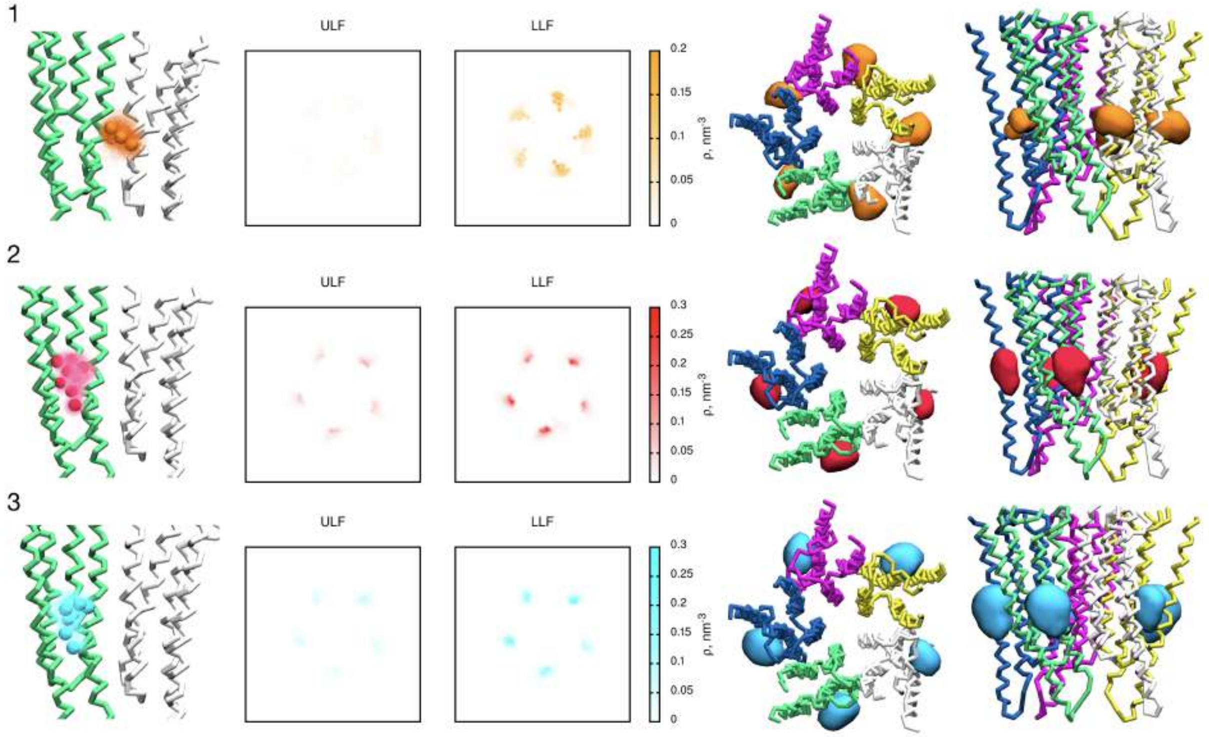

Supplementary Figure 16: Analysis of the most relevant THC binding modes extracted from the CG/MD simulations (see Fig. 2 in *Main Text*). Protein is shown in cartoon representation, in which different colors highlight a different receptor subunit. THC structures, 2D and 3D density maps are colored accordingly to the color code of Fig. 2 in *Main Text*. From the left to the right:
i) binding mode number, as reported in Fig. 2 in *Main Text*;
ii) graphical CG representation of the binding mode degeneracy (as described in Methods. THC is represented as colored beads as the principal binding mode, while all of the other degenerate binding modes as colored overimposed spots;
iii) 2D-density maps in $nm^{-3}$. ULF and LLF define membrane upper and lower, respectively. See text for further details;
iv) 3D-density maps (or volumetric maps) from top and front view of the GlyR. The displayed isosurface corresponds to a density of 0.040 points/Å$^3$. See text for further details.

#### AEA Results

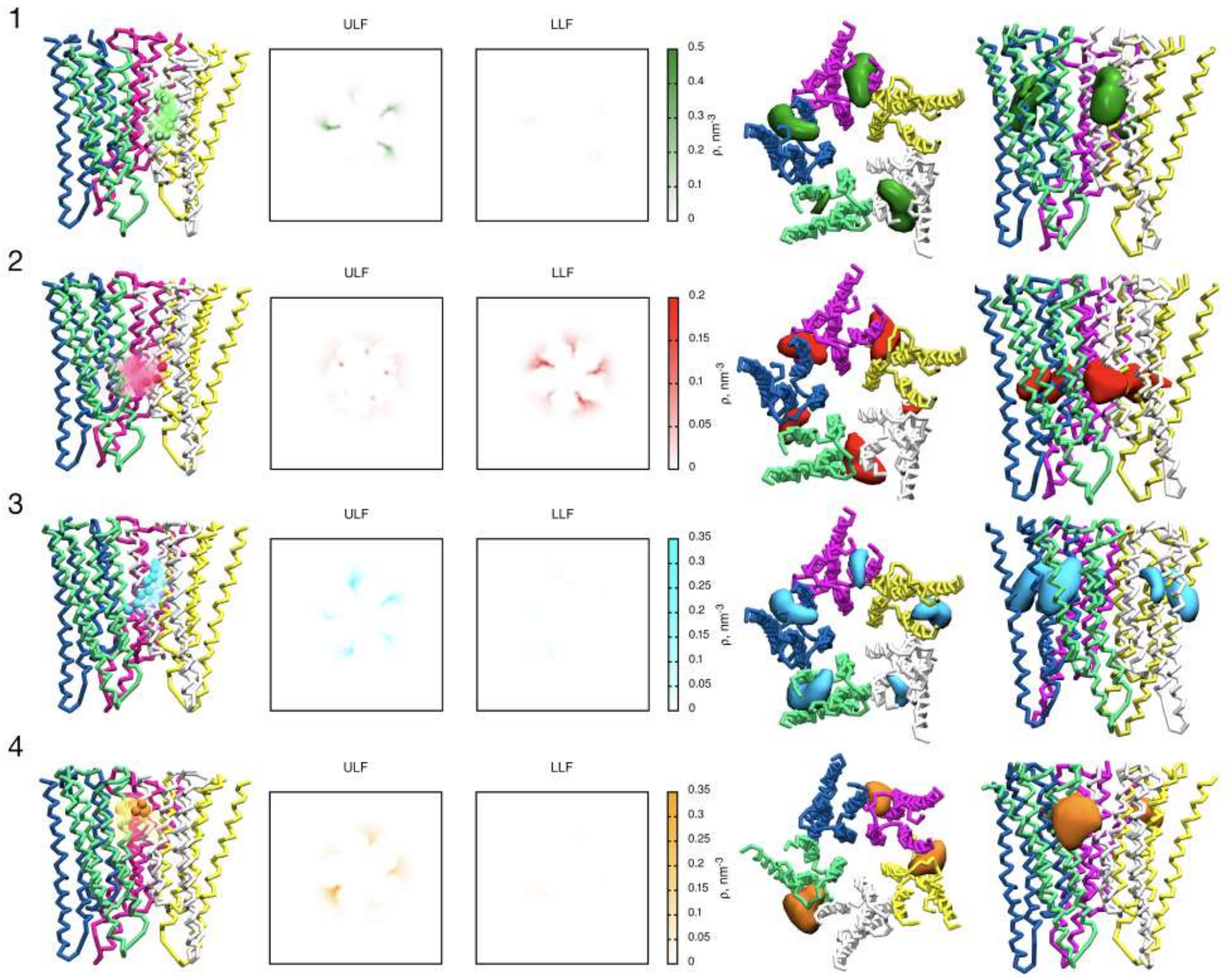

Supplementary Figure 17: Analysis of the most relevant AEA binding modes extracted from the CG/MD simulations (see Fig. 3 in *Main Text*). The protein is shown in cartoon representation, in which different colors highlight a different receptor subunit. AEA structures, 2D and 3D density maps are colored accordingly to the color code of Fig. 3 in *Main Text*. From the left to the right:

i) binding mode number, as reported in Fig. 3 in *Main Text*;

ii) graphical CG representation of the binding mode degeneracy (as described in Methods. AEA is represented as colored beads as the principal binding mode, while all of the other degenerate binding modes as colored overimposed spots;

iii) 2D-density maps in $nm^{-3}$. ULF and LLF define membrane upper and lower, respectively). See text for further details;

iiii) 3D-density maps (or volumetric maps) from top and front view of the GlyR. The displayed isosurface corresponds to a density of 0.040 points/$Å^3$. See text for further details.

#### Water penetration

Water molecules penetrating from the ion pore to the interior of the protein were analyzed over 0.5 ms CG/MD simulation using `gmx densmap` tool. The results in Supplementary Fig. 18 show the existence of significant water channels in the lower leaflet, which produce water density in proximity to the intersubunit cannabinoid-binding site. The presence of a wet protein cavity nearby this site justifies AEA binding with its polar head to an apparent hydrophobic pocket (see *Main Text*).

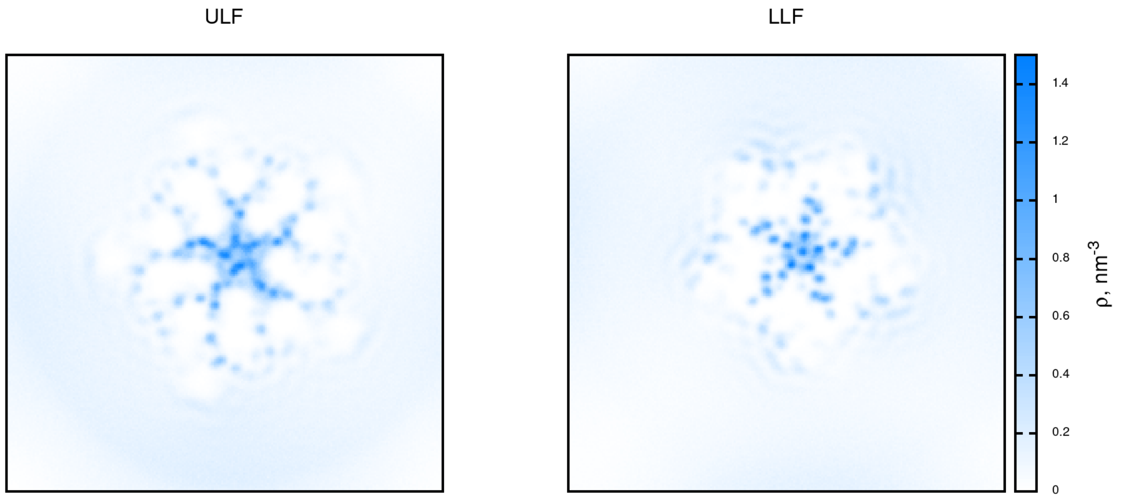

Supplementary Figure 18: 2D-density maps in $nm^{-3}$ of water around GlyR in GlyR-AEA. ULF and LLF define membrane upper and lower, respectively.

#### Protein-Ligand Contact Analysis

By analyzing the concatenated trajectories for the most relevant binding modes, the average number of protein-ligand contacts per residue was obtained. The most important residues for binding THC are shown in Supplementary Fig. 19, those for binding AEA are shown in Fig. 4.

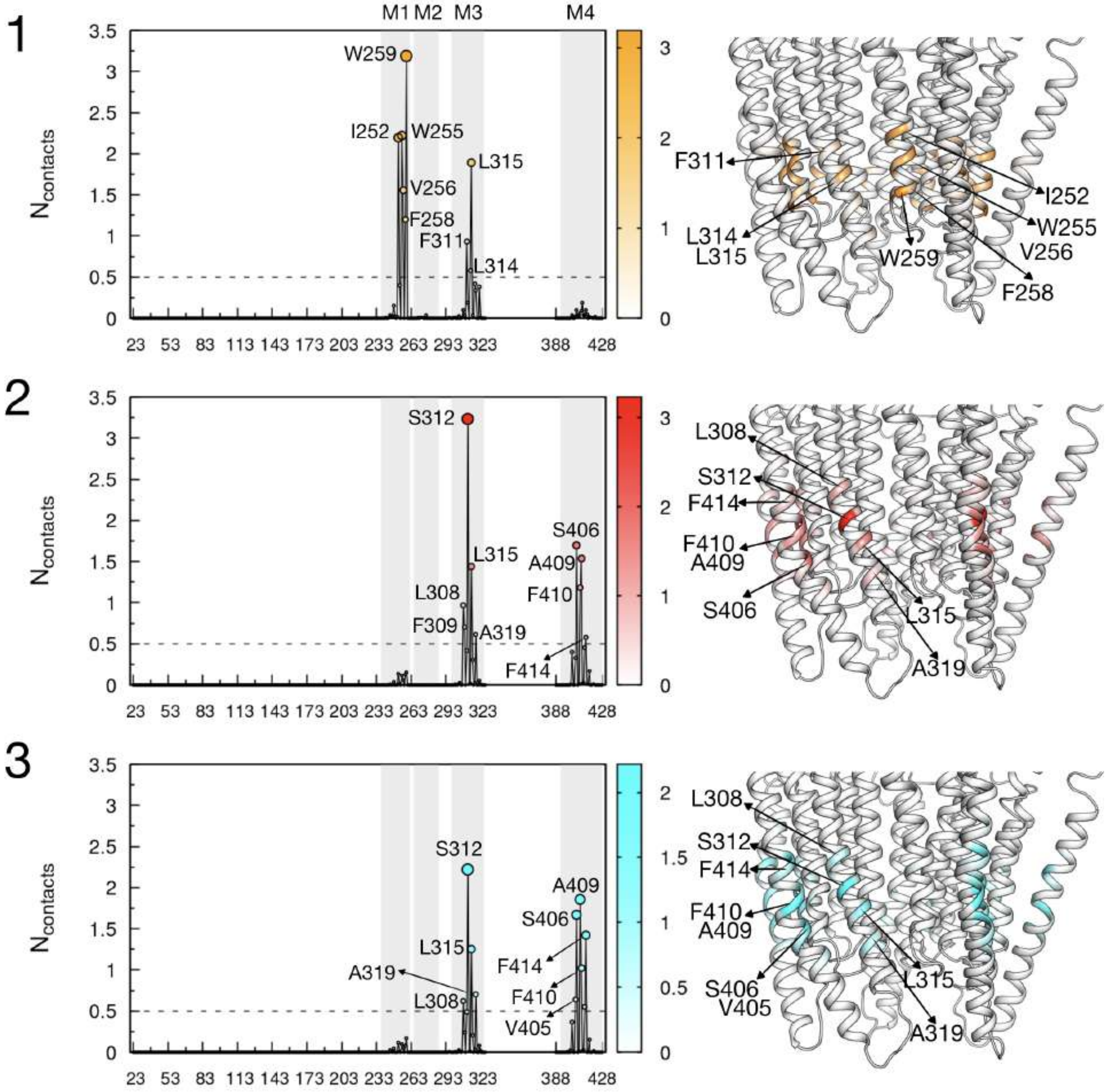

Supplementary Figure 19: Contact analysis carried out for the three specific GlyR-THC binding modes (reported in Supplementary Fig. 16 and Fig. 3 in *Main Text*). On the left, the THC average contacts are reported in function of GlyR residues. The corresponding 3D-color map is reported on the right. The color code follows the one used in Supplementary Fig. 16 and in Fig. 3 in *Main Text*. Gray regions indicate M1, M2, M3 and M4 helices, as reported in panel 1.

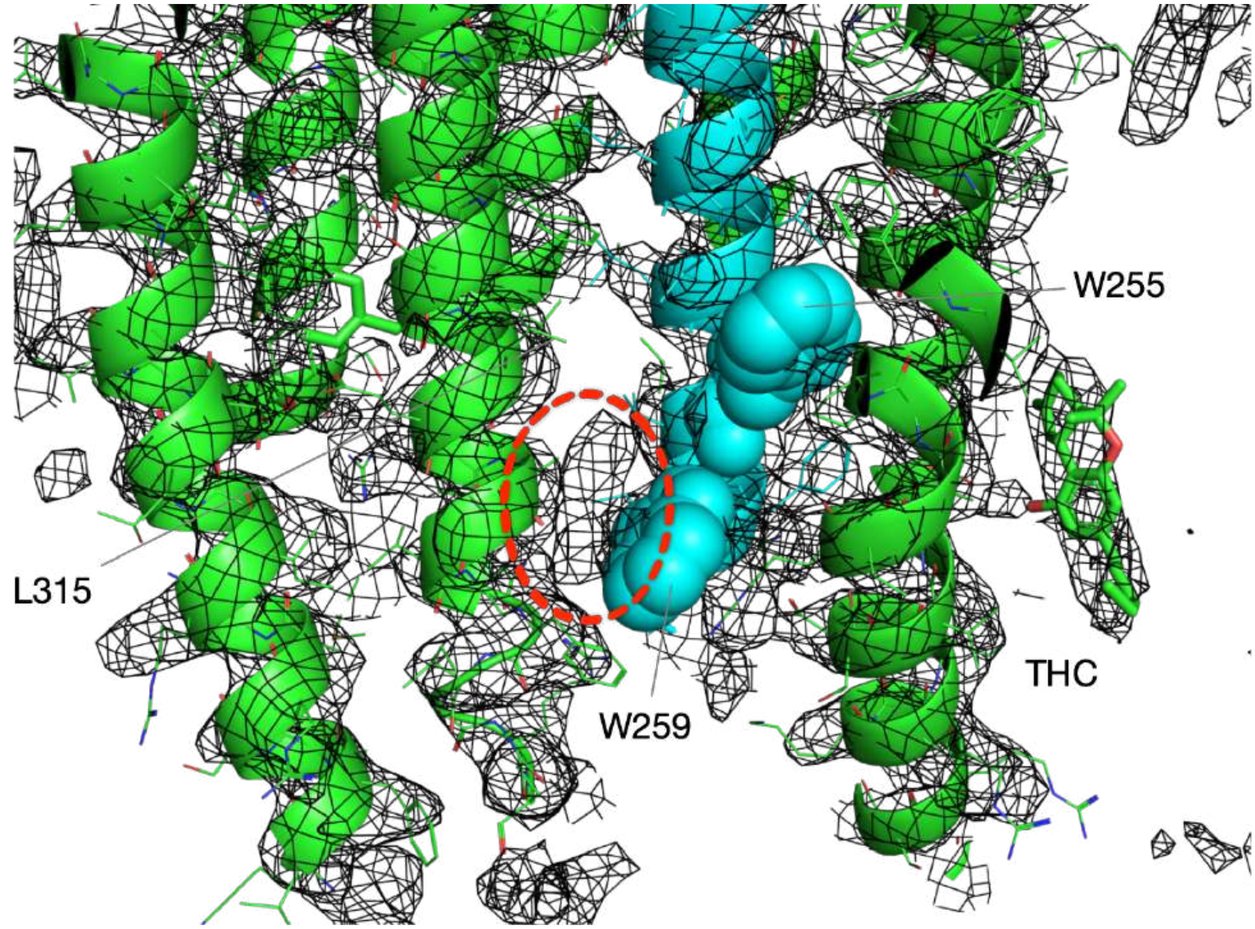

Supplementary Figure 20: The electronic density at one of the recent structures of zebrafish GlyR-$\alpha$1 solved in complex with THC (PDB ID: 7M6M) shows additional un-assigned density consistent with THC intersubunit binding. This additional density (red dashed circle) is located at the interface between subunits, it is compatible with the di-terpenyl core of THC, and is in contact with the bulky side chain of W267 (i.e. W259 in 6PM6). Albeit not being a proof, this evidence supports predictions from the CG/MD simulations.

Supplementary Figure 21: Representative TEVC recordings of GlyR-$\alpha$1 WT and mutants showing $I_{\text{Gly}}$ activated by $EC_{20}$ concentrations of glycine before, during and after co-application with $10\mu M$ AEA in *Xenopus* oocytes expressing GlyR-$\alpha$1 WT or mutants. The membrane potential was hold at $-60$ mV.

A

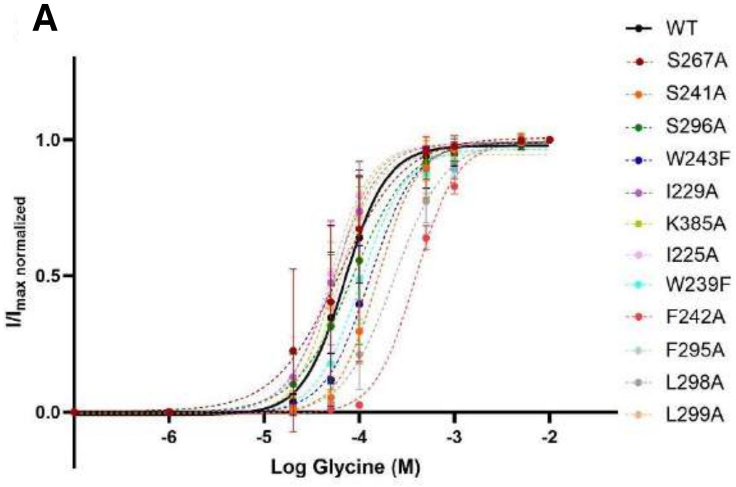

B

| | $EC_{50}$ (μM) | $n_H$ | Evaluated $EC_{20}$ (μM) | n |
|---|---|---|---|---|
| **WT** | 70 ±15 | 1.95 ±0.77 | 28 | n=5 |
| **S267A** | 61 ±23 | 1.28 ±0.68 | 24 | n=5 |
| **S241A** | 157 ±30 | 2.03 ±0.55 | 63 | n=5 |
| **S296A** | 85 ±30 | 1.44 ±0.77 | 34 | n=5 |
| **W243F** | 130 ±28 | 1.82 ±0.82 | 52 | n=5 |
| **I229A** | 54 ±10 | 1.80 ±0.61 | 21 | n=5 |
| **K385A** | 54 ±6 | 2.15 ±0.45 | 21 | n=5 |
| **I225A** | 50 ±7 | 2.05 ±0.56 | 21 | n=5 |
| **W239F** | 101 ±20 | 1.82 ±0.73 | 40 | n=5 |
| **F242A** | 390 ±22 | 2.02 ±0.20 | 155 | n=5 |
| **F295A** | 71 ±23 | 1.99 ±1.37 | 28 | n=5 |
| **L298A** | 231 ±68 | 1.63 ±0.47 | 92 | n=5 |
| **L299A** | 60 ±8 | 2.28 ±0.73 | 24 | n=5 |

Supplementary Figure 22: Glycine dose-response curves. (A) Glycine dose-response curves of GlyR-$\alpha$1 wild-type (solid line) and mutants (dotted line). Data normalized to the maximum current induced by glycine (5mM) with mean ± SD values (n=5). (B) Concentration-response curves parameters (i.e., $EC_{50}$ and Hill coefficient, $n_H$) obtained from the curve fits of normalized concentration-responses to the Hill equation; see Methods.

| Reliability and reproducibility checklist for molecular dynamics simulations | Yes | N/A | Response |
|---|---|---|---|
| **1. Convergence of simulations and analysis** | | | |
| 1a. Is an evaluation presented in the text to show that the property being measured has equilibrated in the simulations (*e.g.* time-course analysis)? | ☒ | | See Fig. 1 and discussion in Results |
| 1b. Then, is it described in the text how simulations are split into equilibration and production runs and how much data were analyzed from production runs? | ☒ | | See "MD simulations" in Methods |
| 1c. Are there at least 3 simulations per simulation condition with statistical analysis? | ☒ | | See "MD simulations" in Methods |
| 1d. Is evidence provided in the text that the simulation results presented are independent of initial configuration? | ☒ | | Production runs of 10 μs can be safely considered as independent |
| **2. Connection to experiments** | | | |
| 2a. Are calculations provided that can connect to experiments (*e.g.* loss or gain in function from mutagenesis, binding assays, NMR chemical shifts, J-couplings, SAXS curves, interaction distances or FRET distances, structure factors, diffusion coefficients, bulk modulus and other mechanical properties, *etc.*)? | ☒ | | Predicted binding modes (Fig. 2 for THC, Fig. 3 for AEA) can be connected to functional studies coupled to site-directed mutagenesis (see Results). Predicted binding affinities in Table 1 could be compared to experiments. |
| **3. Method choice** | | | |
| 3a. Is it described in the text what force field and water model are used and why? | ☒ | | See "Setup for the CG/MD simulations" in Methods |
| 3b. Do simulations contain membranes, membrane proteins, intrinsically disordered proteins, glycans, nucleic acids, polymers, or cryptic ligand binding? | ☒ | ☐ | See "Setup for the CG/MD simulations" in Methods |
| If 3b is **YES**, are enhanced sampling methods used? | ☐ | ☒ | Response not needed if **N/A** |
| **4. Code and reproducibility** | | | |
| 4a. Is a table provided describing the system setup, such as simulation box dimensions, total number of atoms, total number of water molecules, salt concentration, lipid composition (number of molecules and type)? | ☒ | | See "Setup for the CG/MD simulations" in Methods |
| 4b. Is it described in the text what simulation and analysis software and which versions are used? | ☒ | | GROMACS (Methods) alchemical_analysis.py (Supplementary Method 2) VOSSVOLVOX (Supplementary Method 3) |
| 4c. Are initial coordinate and simulation input files and a coordinate file of the final output provided as supplementary files or in a public repository? | ☒ | | See "Data Availability" |
| 4d. Is there custom code or custom force field parameters? | ☒ | ☐ | Response not needed if **N/A** |
| If **YES**, are they provided as supplementary profiles or in a public repository? | ☒ | | CG parameters for THC and AEA are freely accessible from the MAD webserver (see "Data Availability") |

Supplementary Figure 23: Molecular Dynamics simulations checklist. This table explicitly directs readers to information regarding reproducibility of the MD simulations.

(8) Bernazzani, L.; Cabani, S.; Conti, G.; Mollica, V. Thermodynamic study of the partitioning of organic compounds between water and octan-1-ol. Effects of water as co-

solvent in the organic phase. *Journal of the Chemical Society, Faraday Transactions* **1995**, *91*, 649–655.

(17) McQuarrie, D. A. *Statistical Mechanics*; Harper & Row, New York, 1976.

(18) De Jong, D. H.; Schäfer, L. V.; De Vries, A. H.; Marrink, S. J.; Berendsen, H. J.; Grubmüller, H. Determining equilibrium constants for dimerization reactions from molecular dynamics simulations. *Journal of computational chemistry* **2011**, *32*, 1919–1928.

(19) Voss, N.; Gerstein, M.; Steitz, T.; Moore, P. The geometry of the ribosomal polypeptide exit tunnel. *Journal of molecular biology* **2006**, *360*, 893–906.

(20) Humphrey, W.; Dalke, A.; Schulten, K. VMD – Visual Molecular Dynamics. *Journal of Molecular Graphics* **1996**, *14*, 33–38.
